# Individual differences in information processing during sleep and wake predict sleep-based memory consolidation of complex rules

**DOI:** 10.1101/2022.08.23.505024

**Authors:** Madison Richter, Zachariah R. Cross, Ina Bornkessel-Schlesewsky

## Abstract

Memory is critical for many cognitive functions, from remembering facts, to learning complex environmental rules. While memory encoding occurs during wake, memory consolidation is associated with sleep-related neural activity. Further, research suggests that individual differences in alpha frequency during wake (∼ 7 – 13 Hz) modulate memory processes, with higher individual alpha frequency (IAF) associated with greater memory performance. However, the relationship between wake-related EEG individual differences, such as IAF, and sleep-related neural correlates of memory consolidation has been largely unexplored, particularly in a complex rule-based memory context. Here, we aimed to investigate whether wake-derived IAF and sleep neurophysiology interact to influence rule learning in a sample of 35 healthy adults (16 males; mean age = 25.4, range: 18 – 40). Participants learned rules of a modified miniature language prior to either 8hrs of sleep or wake, after which they were tested on their knowledge of the rules in a grammaticality judgement task. Results indicate that sleep neurophysiology and wake-derived IAF do not interact but modulate memory for complex linguistic rules separately. Phase-amplitude coupling between slow oscillations and spindles during non-rapid eye-movement (NREM) sleep also promoted memory for rules that were analogous to the canonical English word order. As an exploratory analysis, we found that rapid eye-movement (REM) sleep theta power at posterior regions interacts with IAF to predict rule learning and proportion of time in REM sleep predicts rule learning differentially depending on grammatical rule type. Taken together, the current study provides behavioural and electrophysiological evidence for a complex role of NREM and REM sleep neurophysiology and wake-derived IAF in the consolidation of rule-based information.

## 1 INTRODUCTION

Rule learning involves forming overarching rules via the extraction of regularities from stimuli that share similar features (Frank & Gibson, 2011). For example, grammar within natural language requires rule extraction, as critical features of a sentence allow for effective categorisation and extraction of meaning (Pothos, 2007). Despite important theoretical and practical implications for understanding the consolidation of rule-based information, the underlying mechanisms that facilitate rule consolidation are poorly understood. Current understanding of rule extraction implicates sleep as an important factor (Batterink et al., 2014; Nieuwenhuis et al., 2013) as well as individual differences in baseline memory performance (Diekelmann et al., 2010). However, individual differences in neurophysiology (cf. Cross et al., 2020a; Klimesch, 1999) as related to rule extraction have yet to be explored in detail.

### 1.1 Memory consolidation as repeated reactivation

The consolidation and generalisation of memory occurs via repeated reactivation of new and existing memory traces (Yassa & Reagh, 2013). Generalisation refers to the process by which the brain extracts and consolidates the central features amongst multiple memory traces, often losing rich contextual details (Stark et al., 2008; Stark & Stark, 2016) and is associated with activity within hippocampus and neocortex (García-Lázaro et al., 2012; Yassa & Reagh, 2013). During generalisation, only central features that overlap between the new memory trace and an existing memory trace are strengthened and the specific, contextual details (e.g., features that do not overlap with the original memory trace), compete for representation in neocortex and are less likely to be consolidated due to mutual inhibition (Yassa & Reagh, 2013). Therefore, over repeated reactivations of similar regularities (e.g., grammar within a language), memory traces become highly semanticised, containing important facts or rules that can be applied to future stimuli (Yassa & Reagh, 2013). This process of consolidation can occur during both periods of wake and sleep (Humiston et al., 2019; Klinzing et al., 2019; Tucker et al., 2020; Wamsley, 2019). In particular, the systems consolidation perspective proposes that memory consolidation and reactivation relies upon slow-wave sleep (SWS; Diekelmann & Born, 2010; Lewis & Durrant, 2011).

### 1.2 The influence of sleep on memory and rule extraction: A systems consolidation perspective

It is now well established that sleep – particularly activity during slow-wave sleep (SWS) – benefits memory consolidation (Diekelmann et al., 2010; Diekelmann & Born, 2010; Lewis & Durrant, 2011; Nieuwenhuis et al., 2013; Stickgold & Walker, 2013; Tononi & Cirelli, 2006). According to the Active Systems Consolidation model (ASC; Diekelmann & Born, 2010), slow oscillations (SOs), thalamo-cortical spindles, and hippocampal ripples facilitate long-term memory consolidation, allowing for newly encoded memories to undergo qualitative changes (e.g., rule extraction) as they are gradually incorporated into existing neocortical memory networks (Klinzing et al., 2019; Lewis & Durrant, 2011). Experimental evidence (Helfrich et al., 2018, 2019; Staresina et al., 2015) demonstrates that phase-amplitude coupling (PAC; Canolty & Knight, 2010) between SOs and faster oscillatory activity (e.g., spindles and hippocampal ripples) facilitates the reactivation of neural assemblies that were active during initial encoding (Diekelmann & Born, 2010; Helfrich et al., 2019; Muehlroth et al., 2019). Additionally, both thalamo-cortical spindles and hippocampal ripples are implicated in the controlled transfer of information, by priming cortical storage areas and modulating selective activation of neural assemblies, respectively (Girardeau & Zugaro, 2011; Staresina et al., 2015). ASC proposes that the PAC between SO and faster oscillatory activity drives the controlled hippocampal-neocortical information transfer required for memory consolidation (Diekelmann & Born, 2010) and more specifically, rule extraction.

Studies exploring rule extraction suggest that sleep facilitates the extraction of overarching rules from regularities within stimuli (Batterink et al., 2014; Djonlagic et al., 2009; Durrant et al., 2011; Gómez et al., 2006; Lerner & Gluck, 2019; Lutz et al., 2017; Nieuwenhuis et al., 2013; Wilhelm et al., 2013), such as grammar within language. For example, a recent review by Lerner & Gluck, (2019) demonstrated that NREM sleep is most associated with the extraction of regularities, however this association is often dependent on the particular task used and whether the regularity contains temporal aspects. Experimental studies investigating the effect of sleep on linguistic rule learning found that greater SWS duration predicted subsequent sensitivity to a hidden rule, as well as being instrumental in stabilising memory traces related to a rule (Batterink et al., 2014; Nieuwenhuis et al., 2013). Additionally, studies in non-linguistic contexts have reported similar results, in which SWS duration was positively associated with the extraction of statistical rules and subsequent performance on a task in which understanding of the rule was crucial (Djonlagic et al., 2009; Durrant et al., 2011, 2013). However, only a few experiments have explored how precise oscillatory events (e.g., SO-spindle coupling) that occur during SWS may predict rule extraction. The limited work in this area has found that stronger SO activity is positively associated with motor sequence learning in both children and adults (Wilhelm et al., 2013), that SO-spindle coupling increases from childhood to adolescence, and this increase predicts improvements in motor memory (Hahn et al., 2020). Beyond motor memory, other research has explored the role of SOs, spindles and their coupling in other memory contexts such as associative memory outcomes across the lifespan (Muehlroth et al., 2019) and declarative memory recall (Mikutta et al., 2019), providing evidence for NREM oscillatory events in facilitating memory consolidation. Despite this, little is known regarding the influence of SO-spindle coupling in other contexts, such as complex rule learning. Due to SO-spindle coupling being linked with general memory consolidation, further investigation of such activity in a rule-based context may increase our understanding of the underlying mechanisms involved in rule extraction.

### 1.3 Beyond slow-wave sleep: A possible role for REM sleep in rule generalisation

In addition to cortical activity within SWS, Rapid Eye Movement (REM) sleep and its associated oscillatory activity has often been implicated in memory consolidation (Ackermann & Rasch, 2014; Boyce et al., 2016; Grosmark et al., 2012). Few studies have investigated the role of neurophysiology during REM (e.g., theta activity; ∼4–7 Hz) within rule-based contexts, typically focussing on the influence of REM sleep duration on processes such as gist extraction (Batterink et al., 2014; Matorina & Poppenk, 2019), which refers to the extraction of the most important elements or the general idea (Matorina & Poppenk, 2021). For example, Matorina & Poppenk (2019) found that a higher proportion of time spent in REM sleep led to decreased performance in a statistical learning task, suggesting that REM sleep is disadvantageous to the extraction of rules. In contrast, Batterink et al., (2014) examined how NREM and REM sleep duration during a nap contributed to rule generalisation and found that performance was positively associated with the interaction between proportion of time spent in REM and NREM sleep, indicating a role for REM sleep in rule extraction.

Other studies have indicated a beneficial effect of REM sleep on rule extraction, specifically when looking at acquisition of a rule for effective categorisation (Djonlagic et al., 2009) and within a task in which effective learning of sequential rules was critical (Maquet et al., 2000). However, research into the specific influence of neurophysiology that occurs during REM sleep on rule extraction is still limited, requiring further exploration. Interestingly, work into the facilitation of memory generalisation has implicated factors other than specific sleep neurophysiology, such as individual differences in memory performance (Diekelmann et al., 2010). Despite this, individual differences in neural activity and their relation to sleep-associated memory consolidation are only just starting to be considered (Chatburn et al., 2021; Cross et al., 2020a; Fenn & Hambrick, 2012; Schabus et al., 2008).

### 1.4 Individual alpha frequency as a measure of individual differences in information processing

Research into inter-individual differences in cognitive processing have often focussed on alpha activity during wake (∼8-12Hz) and its relation to information processing in a variety of domains, such as language and memory (Bornkessel et al., 2004; Bornkessel-Schlesewsky et al., 2022; Cross, Santamaria, et al., 2020; Furman et al., 2018; Klimesch, 2012; Samaha & Postle, 2015). The individual alpha frequency (IAF), measured during wake, refers to the individual dominant peak within the alpha frequency band, and has been linked to various aspects of cognition such as attention, processing speed and memory (Klimesch, 1997, 1999; Surwillo, 1961). IAF has been implicated in the controlled timing of information transmission (Klimesch, 2012) and it is hypothesised that those with a higher IAF have improved information transmission within cortex, resulting in better performance on tests of memory (Klimesch, 1999) and faster processing speed (Klimesch et al., 1996; Ociepka et al., 2022; Surwillo, 1961, 1963). Samaha and Postle (2015) suggest that IAF during wake may be related to temporal receptive windows of processing, determining the amount of information within each processing cycle. Here, higher IAF individuals are considered to have faster cycles with less information contained within, and lower IAF individuals have slower cycles that contain more information (Samaha & Postle, 2015). In relation to memory, research indicates that those with good memory performance tend to have an IAF of around 1 Hz higher than poor memory performers (Doppelmayr et al., 2005; Klimesch, 1997, 1999).

The proposed role of IAF in processing speed and information transfer described above may have implications for processes dependent on information processing and cortical communication. Therefore, as sleep-dependent memory consolidation is proposed to rely heavily on information transmission within the brain, mechanisms suggested to reflect this process (e.g., SO-spindle coupling) may interact with IAF due to their shared association with information processing and transfer. Whilst there is little experimental evidence for an explicit association between sleep neurophysiology and wake-state IAF, Cross et al., (2020a) investigated how these factors may interact to influence emotional memory consolidation. Results revealed complex interactions between wake-derived IAF, REM theta power and SO density modulating emotional memory outcomes, indicating a relationship between individual processing capabilities during wake and cortical activity during sleep, providing initial support for a link between IAF and sleep neurophysiology. However, the interaction between these variables in other memory-related contexts (e.g., rule extraction) is yet to be examined.

### 1.5 The current study

The present study is a reanalysis of Cross et al. (2021) and aimed to address whether sleep-related (e.g., SO-spindle coupling) and intrinsic neural differences in information processing (e.g., IAF) modulate rule extraction. Due to the significance of individual differences in memory performance, exploring IAF as a measure of individual differences will inform current models of sleep and memory as to how underlying neurophysiology of an individual interacts with memory consolidation and thus, rule extraction. Exploratory analyses further aimed to clarify the role of REM sleep and its involvement in rule-based memory consolidation. Subjects were randomly allocated to a sleep or wake condition and were required to learn grammatical rules in a novel artificial language modelled on Mandarin Chinese (Cross et al., 2020b). Participants then completed a baseline and delayed sentence judgment task (after sleep or an equivalent period of wake) used to test knowledge about underlying grammatical rules, performance on which served as a proxy for rule extraction.

Based on previous literature suggesting the role of sleep in facilitating memory performance, it was hypothesised that (H^1^) those in the sleep condition will perform better on tasks assessing grammatical knowledge than those in the wake condition. Additionally, due to previous literature indicating a beneficial effect of sleep and IAF on memory performance, it was hypothesised that (H^2^) differences in memory performance relative to condition (sleep vs. wake) and grammatical rule type will be modulated by both SO-spindle coupling and IAF and that this relationship may be (a) additive or (b) interactive. Additionally, exploratory analyses were conducted to assess whether REM sleep is associated with the extraction of complex rules.

## 2 METHODS

### 2.1 Participants

Thirty-five (16 males) adults ranging from 18 to 40 years of age (*M* = 25.4, *SD* = 7.10) were randomly allocated to either a sleep (*Mean age* = 23.60 [*SD* = 5.84], 9 males) or wake (*Mean age* = 27.11 [*SD* = 7.50], 8 males) condition (17 in sleep condition). Eligible participants were healthy, right-handed, had no current or past psychiatric conditions (including intellectual impairment) or history of substance dependence, normal or corrected-to-normal vision and were not taking medication that influenced neurophysiological measures. Participants were instructed to not consume caffeine prior to the experiment. In addition, eligible participants were monolingual, native English speakers with no previous exposure to Mandarin Chinese and did not have any current conditions that affected sleep or circadian rhythms and had not undertaken shift work in the past month to ensure no influence of irregular or poor-quality sleep. Participants received an AUD$120 honorarium upon completion of the study. Ethics approval was granted by the University of South Australia’s Human Research Ethics committee (I.D: 0000032556).

### 2.2 Materials and Measures

#### 2.2.1 Electroencephalography (EEG)

The EEG was recorded with a 32-channel BrainCap (Brain Products GmbH, Gilching, Germany) with Ag/AgCl electrodes mounted via the International 10-20 system. The ground electrode was located at AFz and the online reference was located at FCz, with re-referencing to linked mastoids undertaken offline. The electrooculogram (EOG) was recorded using electrodes located at the side of each eye as well as above and below the left eye. All EEG and EOG channels were amplified via a BrainAmp DC amplifier (Brain Products GmbH, Gilching, Germany), band-pass filter of DC – 250Hz and a sampling rate of 1000 Hz. Impedances were kept below 10kΩ. EEG signals were recorded throughout both the sleep durations and experimental tasks.

#### 2.2.2 Language Learning Task: Mini Pinyin

Mini Pinyin is a modified miniature language (MML) modelled on Mandarin Chinese (Cross et al., 2020b). MML paradigms contain words that are combined into sentences following grammatical rules adapted from the language they are based upon (Mueller, 2006; Wang et al., 2012). Mini Pinyin contains 16 verbs, 25 noun phrases, 2 coverbs, and 4 classifiers (for a full description of Mini Pinyin, see Cross et al., 2020b). The nouns are divided into categories of human (10 nouns), animal (10 nouns), and objects (5 nouns).

Mini Pinyin contains two coverbs that influence word order and the interpretation of the noun phrases. For example, when bǎ is present in a sentence, the first noun is the Actor (i.e., the participant instigating the event) and the second noun is the Undergoer (i.e., the participant affected by the event). The second coverb, bèi, reverses this pattern. When either of these coverbs are used, word order becomes fixed and determines role assignment (see Table 1 for examples). Thus, bǎ constructions instantiate Actor-first word orders, while bèi instantiates an Undergoer-first word order. Additionally, verb positioning is an important element in Mini Pinyin: sentences that contain coverbs must have the verb placed at the end of the sentence, whereas sentences without coverbs (not examined in this analysis) require the verb to be positioned between the noun phrases.

**Table 1.** Examples of plausible and implausible sentence structures (adapted from Cross et al., 2020b).

| Coverb | Sentence Examples |
| --- | --- |
| *Bǎ* Plausible (Actor-first) | zhi junma bǎ xi pingguo chile.<br>(animal) horse bǎ (small object) apple eat.<br>*"The horse eats the small apple."* |
| *Bǎ* Implausible (Actor-first) | xi pingguo bǎ zhi junma chile.<br>(small object) apple bǎ (animal) horse eat.<br>*"The small apple eats the horse."* |
| *Bèi* Plausible (Undergoer-first) | da shubao bèi ge yisheng dale.<br>(large object) bag bèi (human) doctor hit.<br>*"The doctor hits the large bag."* |
| *Bèi* Implausible (Undergoer-first) | ge yisheng bèi da shubao dale.<br>(human) doctor bèi (large object) bag hit.<br>*"The large bag hits the doctor."* |

Based on the linguistic parameters described above, the process of rule extraction can be explored by using the word order contingencies (e.g., the fixed word order of the Actor-Undergoer relationship) of each of the coverbs. Each coverb and the associated rules can be used to examine rule extraction across differing levels of novelty in comparison to English (i.e., the native language of the participants). While both coverbs require participants to learn a novel verb positioning rule in comparison to English, Actor-first bǎ-constructions show some similarity to canonical English word order in that the Actor precedes the Undergoer. Undergoer-first bèi-constructions, by contrast, differ from English word order in two respects: the position of the verb and the relative order of the nouns. Accurate rule extraction would allow the participant to correctly distinguish between the specific noun phrase order instantiated by Undergoer-first (bèi) and Actor-first (bǎ) constructions as well as the correct position of the verb. Table 2 summarises the degree of novelty of each construction in comparison to English.

**Table 2.** Summary of similarities (+) and differences (-) to English word order for Actor-first (Bǎ) and Undergoer-first (Bèi) coverb constructions within Mini Pinyin.

| Sentence Type | Noun Order | Verb Position |
| --- | --- | --- |
| *Bǎ* (Actor-first) | + | - |
| *Bèi* (Undergoer-first) | - | - |

**Note:** Other sentence types in Mini Pinyin (not examined in this analysis) required the verb to be positioned similarly to English. As such, participants were required to learn the verb-final positioning rule that was unique to the sentences with coverbs.

### 2.3 Procedure

Each participant received a picture-word pair booklet containing each of the 25 nouns approximately one week before testing to gain a basic vocabulary before the experimental tasks. Eligible participants visited the Cognitive Neuroscience Laboratory at the University of South Australia to undergo testing. Before continuing onto the main testing phase, participants were required to score above 90% on a vocabulary test designed to test their knowledge of the 25 nouns. All 36 participants included in this analysis scored above 90%.

The main testing session consisted of a sentence learning task, a baseline sentence judgement task, and a delayed judgement task (approximately 12 hours after the learning and baseline tasks). All sentence and picture stimuli were presented via OpenSesame (Mathôt et al., 2012). The learning task used pictures to depict events involving two entities and was designed for the participants to learn the meaning of the verbs, coverbs, classifiers, and overall sentence structure. Each trial began with a 1000ms fixation cross, followed by pictures depicting the event. Subsequently, a sentence describing the event pictured was presented on a word-by-word basis, with each word presented for 700ms. The inter-stimulus interval (ISI) was 200ms. This trial structure was used for a total of 128 pseudorandomised picture-sentence combinations.

Following the learning task, participants completed the baseline and delayed judgement tasks. Both judgment tasks consisted of 288 pseudorandomised novel sentences (without pictures) presented word-by-word, with each word being presented for 600ms each with an ISI of 200ms. Participants were instructed prior to the task to judge if the sentence presented was grammatically correct by pressing a button (cued by a question mark presented on screen for 4000ms). During the baseline judgment task, participants also received feedback as to whether their response was correct. They did not receive feedback in the delayed judgement task. Two stimuli lists were used to counterbalance the order of stimuli presentation across participants. Half of the sentences were grammatical constructions, and the other half were ungrammatical constructions. See Figure 1 for a schematic of both the learning and baseline judgement tasks.

**Figure 1.**
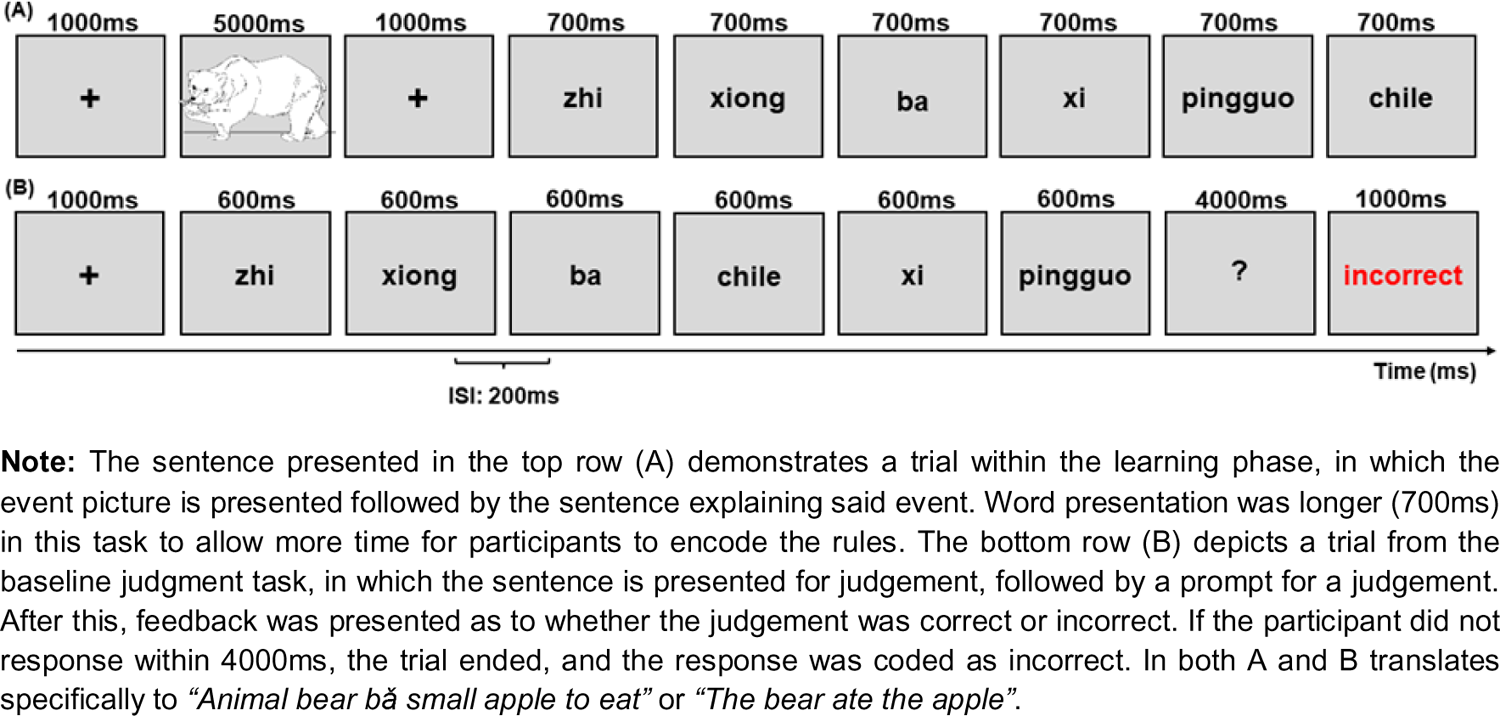
Schematic representation of trials within the (A) learning and (B) baseline judgement tasks (adapted from Cross et al., 2021).

Participants in the wake condition completed the learning and baseline judgment tasks at approximately 09:00hr before leaving the laboratory to undergo their normal daily routine. They were instructed not to nap or engage in any language learning activities that may interfere with the task. Participants returned the same day to complete the delayed judgment task at approximately 21:00hr.

Participants in the sleep condition completed the learning and baseline judgment tasks at approximately 21:00hr after EEG set up. Following this, participants were given an 8-hour sleep opportunity between 23:00hr – 07:00hr. For the sleep opportunity, EMG was recorded, EOG channels were removed from the vertical position (above and below left eye) and placed horizontally (either side of each eye), and polysomnography was continuously recorded. Upon waking, participants were given approximately 1 hour to overcome sleep inertia and 1 hour for EEG and poor impedances to be fixed before completing the delayed judgement task (at approximately 09:00hr). For a schematic of the experimental protocol refer to Figure 2.

**Figure 2.**
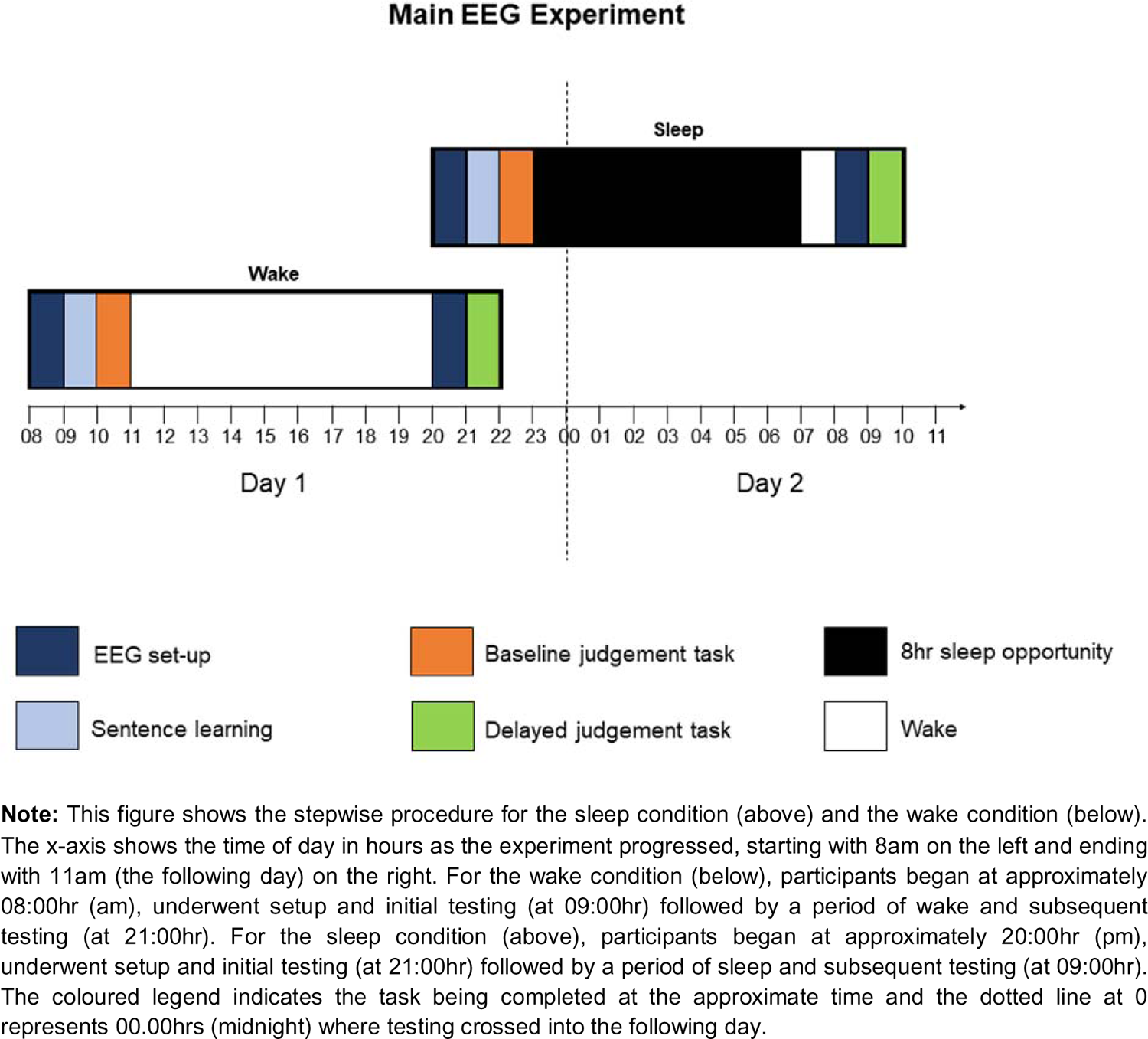
Schematic representation of experimental protocol (adapted from Cross et al. 2021).

### 2.4 Data Analysis

All statistical analyses were performed in *R* (R Core Team, 2017). Mixed-effects regression models were fitted using maximum likelihood via the *lme4* (Bates et al., 2015). Other packages included in mixed-model analysis and output reporting included the *car* (Fox & Weisberg, 2018), *Rmisc* (Hope, 2013), *effects* (Fox & Weisberg, 2018), *Hmisc* (Harrell Jr, 2019), *apaTables* (Stanley & Spence, 2018) and *performance* (Lüdecke et al., 2021). Data was arranged using *tidyverse* (Wickham et al., 2019), *reshape2* (Wickham, 2007), *plyr* (Wickham, 2011), *sjmisc* (Lüdecke, 2018a), and packages involved in plotting included *lattice* (Sarkar, 2008), *sjPlot* (Lüdecke, 2018b), *ggpubr* (Kassambara, 2020), *RColorBrewer* (Neuwirth, 2014). All effects were plotted using the *ggplot2* package (Wickham et al., 2021) and *ggeffects* (Lüdecke et al., 2020). Rayleigh’s test for non-uniformity was run using *CircStats* (Lund & Agostinelli, 2012). Links to the raw data can be found in the supplementary material (Appendix E.1) and analysis scripts can be found at the following Open Science Framework repository: https://tinyurl.com/turbidus.

#### 2.4.1 Behavioural analysis

Performance on both the baseline and delayed tasks were quantified using d-prime (Lockhart & Murdock, 1970; Wixted, 2014). d-prime is a measure of discriminability and is defined as the difference between the z-transformed scores of hit rate (HR) and false alarm rate (FA; i.e., d-prime = z[HR] – z[FA]). To calculate d-prime, each sentence type was isolated and scored according to the criteria described in Table 3. For example, calculating d-prime for bǎ verb position sentences consisted of isolating bǎ verb position sentences and scoring them. Here, grammatical bǎ sentences that were correctly identified as grammatical were scored as a hit, whilst ungrammatical bǎ verb position sentences that were incorrectly identified as grammatical were scored as a false alarm. Following this, the number of hits and false alarms per subject were converted to a proportional value before being z-transformed and applying the d-prime calculation. Raw hit and false alarm scores for each participant per sentence type can be found in the supplementary material. For example, for bǎ verb position sentences, a formula of z(HR(bǎ grammatical) – z(FA(bǎ ungrammatical)) was applied. d-prime calculations for the baseline testing session were also calculated to be added into the behavioural linear mixed-effects models for comparison to delayed scores. Baseline scores were also added into subsequent sleep neurophysiology models to control for baseline performance when predicting delayed memory outcomes. Positive d-prime scores indicate good memory performance, whereas negative scores indicate poor memory performance. d-prime scores of 0 reflect chance performance.

**Table 3.**
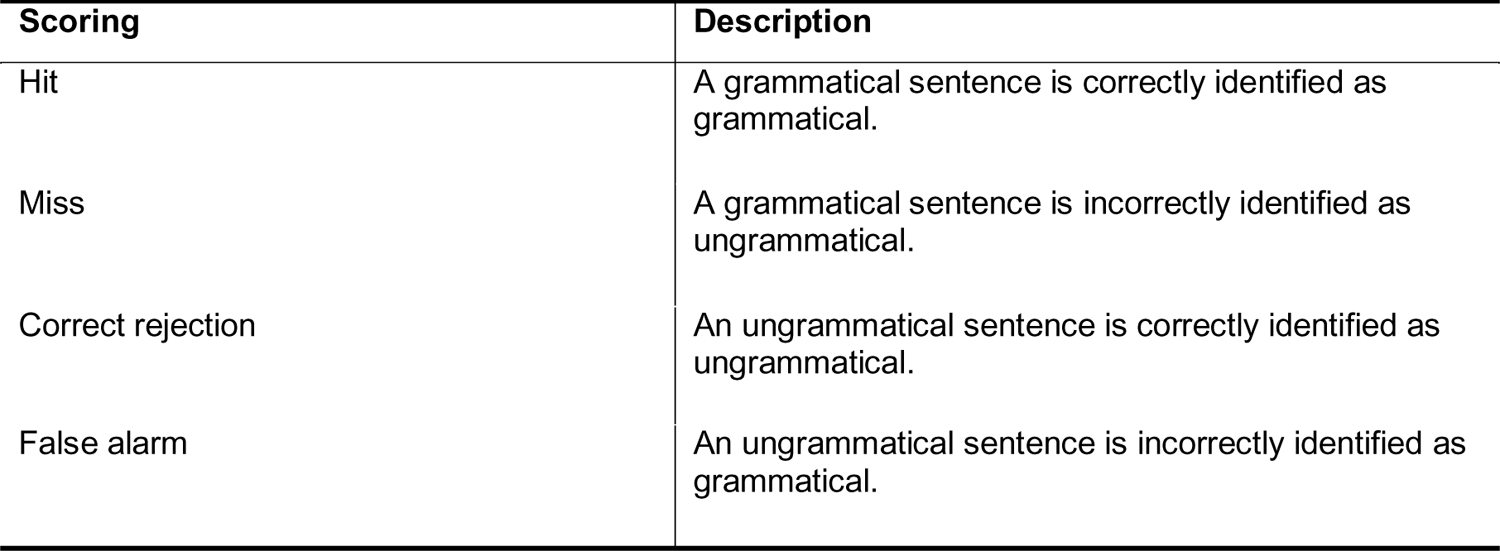
Description of how responses were scored to calculate d-prime for the delayed testing phase for measurement of memory performance.

#### 2.4.2 Sleep EEG analysis

Sleep EEG was pre-processed according to the specifications detailed in (Cross et al., 2021). Briefly, sleep EEG was scored according to standardised criteria (AASM; Berry et al., 2012) by two sleep technicians using Compumedics Profusion 3 software (Melbourne, Australia). A high-pass filter of 0.3 Hz and a low-pass filter of 35 Hz were applied for viewing. For the analysis, the EEG was filtered from 0.1 – 40 Hz and re-referenced to linked mastoids. Artifact rejection was performed automatically, in which a covariance-based artifact rejection method was used with hypnograms (Barachant, 2015; Barthélemy et al., 2019). Additionally, the EEG was down-sampled from 1000 Hz to 100 Hz to reduce computational processing time.

Phase-amplitude coupling (PAC) events were detected using published algorithms (Dvorak & Fenton, 2014; Helfrich et al., 2019; Staresina et al., 2015) via the YASA toolbox implemented in MNE Python (Vallat & Walker, 2021). An event-locked method was utilized, in which the SOs were detected initially, and spindle events that occur within the detected SOs were subsequently identified. The PAC strength between SOs and spindles was calculated using the Mean Vector Length (MVL) method (Canolty et al., 2006). MVL is calculated by combining the amplitude of the frequency for amplitude (i.e. spindle) with the phase of the frequency for phase (i.e. SO), in which the combined signals become complex vectors (Canolty et al., 2006; Hülsemann et al., 2019; Munia & Aviyente, 2019). The average length of these vectors can be delineated, whereby the vector length represents the magnitude (or strength) of the coupling event (Hülsemann et al., 2019).

For the REM theta analysis, power spectral density (PSD) estimates were calculated on consecutive 30s epochs with a 4-second sliding window. Intervals were tapered by a single Hanning window before applying a fast Fourier transformation that resulted in interval power spectra with a frequency resolution of 0.25 Hz. Power spectra were then averaged across all blocks (Welch’s method) to obtain PSD estimates for relative theta power (4 – 8 Hz) during REM epochs for each channel (for a detailed description of this approach, see Vallat & Walker, 2021).

#### 2.4.3 Individual alpha frequency calculation

Individual alpha frequency (IAF) estimates for each subject were calculated from eyes-closed resting-state EEG recordings according to the method detailed by Corcoran et al., (2018), via the *philistine* package (Alday, 2019). Resting-state recordings were taken at the beginning of the learning task and after the judgment task for 2 minutes (eyes open and eyes closed). Estimates were taken using the peak alpha frequency (PAF) method, in which the most prominent peak in the alpha bandwidth was delineated. The PAF for each participant was estimated from occipital and parietal channels (P3/P4/P7/P8/Pz/O1/O2/Oz) as this is where the alpha rhythm is most prominent during resting-state eyes closed periods (Babu Henry Samuel et al., 2018; Capotosto et al., 2017). For a detailed description of the IAF estimation procedure, please see Corcoran et al., (2018).

#### 2.4.4 Statistical analysis

Data were analysed using linear mixed-effects models via the *lme4* package (Bates et al., 2015). For all models, Participant ID was specified as a random effect on the intercept (Brown, 2020) and significance of effects were examined using Type II Wald tests from the *car* package (Fox & Weisberg, 2018).

Model predictors (fixed effects) included Condition (sleep, wake), Coverb (bǎ, bèi), Violation Type (noun phrase order, verb position), PAC events (quantified as MVL), IAF, and Baseline d-prime scores. d-prime scores from the delayed testing session were specified as the outcome variable. Categorical variables were sum-to-zero contrast coded, meaning that factor-level estimates are relative to the grand mean (Schad et al., 2020). The first model was run exploring the effect of Condition (sleep vs. wake), Coverb, Violation Type, baseline d-prime scores, and IAF on memory performance. A second model was run to investigate the effect of PAC, Coverb, Violation Type, baseline d-prime scores and IAF on memory performance to explore neurophysiological factors in detail.

Two additional exploratory models were run to investigate the role of REM sleep and associated theta power on memory performance. Predictors of the initial model included IAF, Coverb, Violation Type, Baseline d-prime scores, REM Theta power and the topographical components saggitality (anterior, central, posterior) and laterality (left, midline, right), to explore oscillatory influences during REM on delayed d-prime scores. An additional model was run with predictors of IAF, Coverb, Violation Type, Baseline d-prime Scores and Percentage of Time in REM Sleep, to investigate how REM sleep influences delayed d-prime scores. REM theta power and percentage of time spent in REM were converted to z-scores.

## 3 RESULTS

### 3.1 Descriptive Statistics

Overall, the average d-prime score across sentences for the wake group were .66 (*SD = .97*) at baseline testing and .65 (*SD = 1.34*) at delayed testing. For the sleep condition, the average d-prime scores across sentence types were .60 (*SD* = .85) at the baseline testing session and .87 (*SD* = 1.40) at the delayed testing session (refer to Table 4 for d-prime scores per sentence type). Raw values for hits and false alarms per subject and sentence condition can be found in the supplementary material (Appendix C). Sleep variables including total sleep time, sleep onset latency, wake after sleep onset, and the total amount of time spent in each sleep stage are reported in Table 4, while Figure 3 shows a comodulogram and circular histogram to illustrate the estimated PAC metrics. Additionally, a correlogram showing pairwise comparison between variables of interest can be found in the supplementary material (Appendix B, Figure 3).

**Figure 3.**
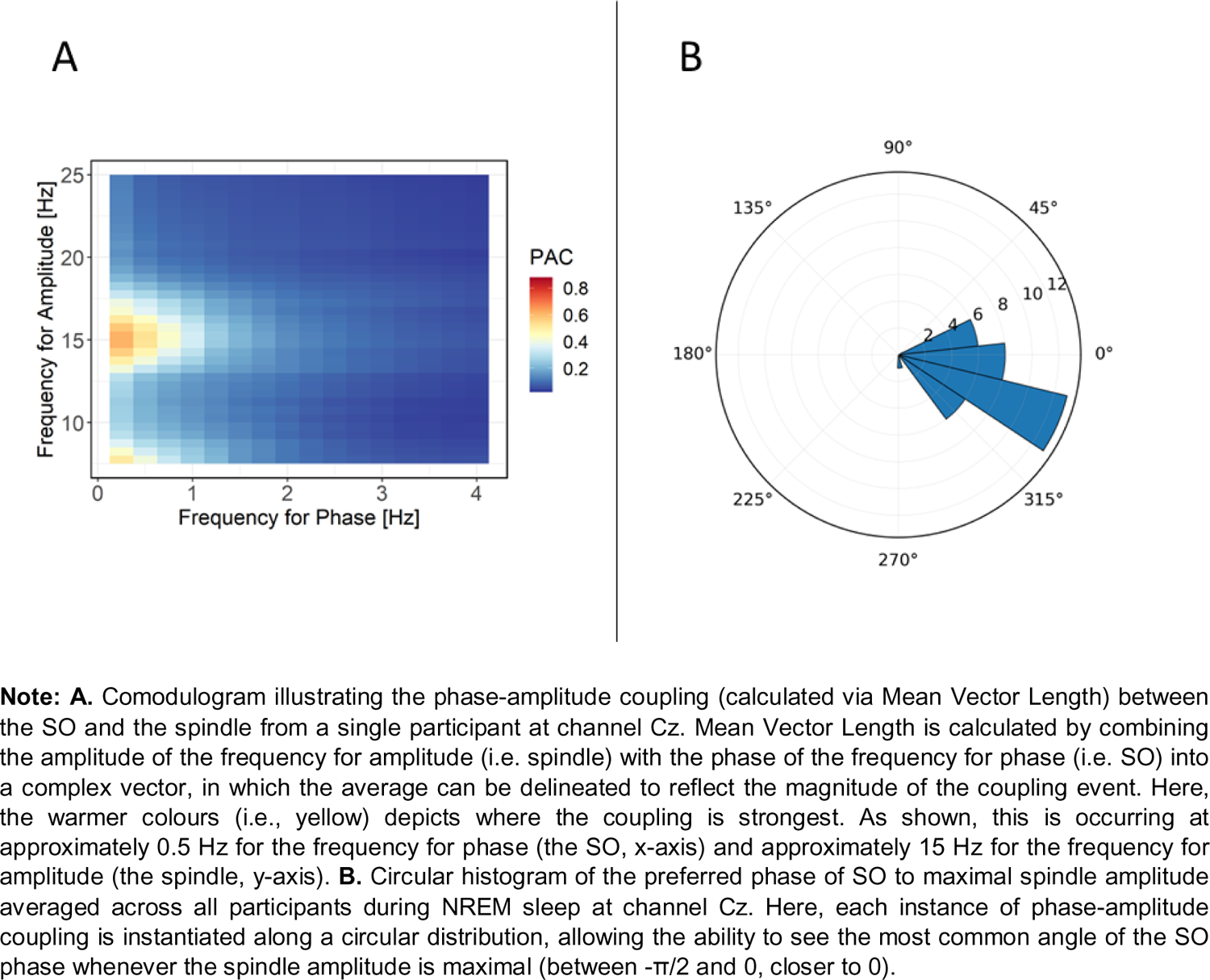
Examples of phase-amplitude coupling metrics for visualisation purposes.

**Table 4.** Mean d-prime scores (and standard deviations) for baseline and delayed testing sessions in the sleep and wake groups for each sentence type.

| Violation Type | Sleep |  | Wake |  |
| --- | --- | --- | --- | --- |
|  | Baseline | Delayed | Baseline | Delayed |
| ***Bă*** |  |  |  |  |
| Noun Order | .26 (.57) | .40 (1.13) | .16 (.52) | .15 (.86) |
| Verb Position | .86 (.80) | 1.37 (1.26) | 1.40 (.91) | 1.20 (1.43) |
| ***Bèi*** |  |  |  |  |
| Noun Order | .29 (.68) | .49 (1.18) | -.01 (.68) | .07 (1.03) |
| Verb Position | .98 (1.08) | 1.25 (1.82) | 1.09 (.94) | 1.21 (1.54) |

**Note:** Standard deviation is in parentheses.

### 3.2 Behavioural Analysis

The results reported in the following summarise the effects most pertinent to the current behavioural hypotheses, drawing on mixed model output summaries (for β values) and Type II Wald Chi-square test for chi-square and *p* values. Full model and Wald test summaries can be found in the supplementary material (Appendix A.1).

Mixed model analyses revealed a significant interaction between Condition, IAF and Baseline d’ predicting d-prime at delayed testing, as shown in Figure 4A (refer to Table 6 for inferential statistical information for all main effects and significant interactions). Overall, a positive relationship between baseline and delayed performance can be seen. However, the strength of this relationship differs with respect to IAF and Condition. In particular, the relationship between baseline and delayed performance is weaker for high IAF individuals in the wake condition, suggesting that wake appears to interfere with rule extraction for high-IAF individuals.

**Figure 4.**
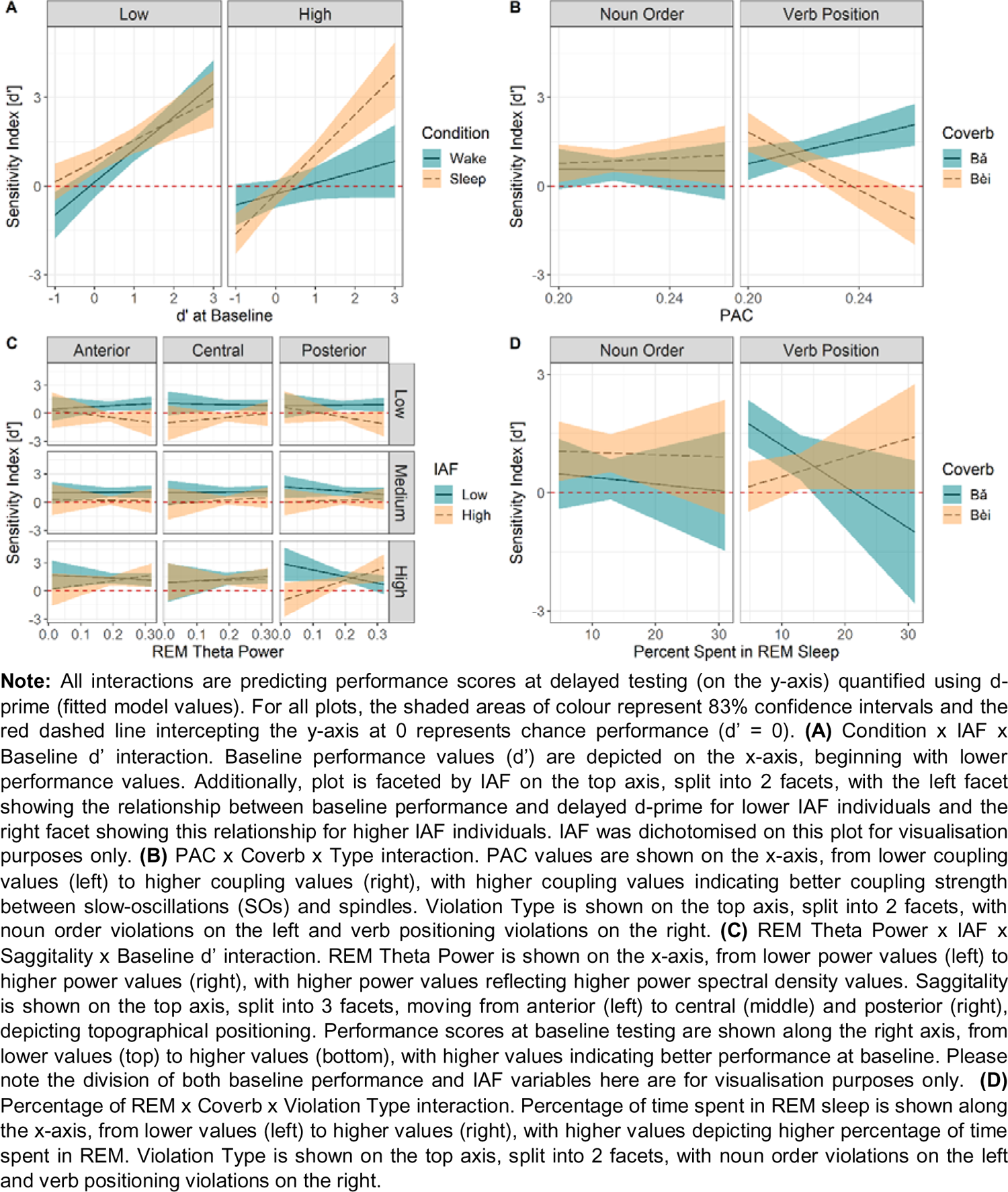
Plotted interactions from the behavioural model (A), sleep model (B), and exploratory models examining REM Theta Power (C) and percentage of time in REM sleep (D).

**Table 5.** Descriptive statistics (means, standard deviations, and ranges) for sleep parameters measured during the 8-hour sleep opportunity between baseline and delayed testing.

| Sleep Stage | Mean Minutes (SD) | % of TST (SD) |
| --- | --- | --- |
| N1 | 38.06 (29.93) | 10.05 (8.34) |
| N2 | 196.29 (47.01) | 49.52 (10.52) |
| SWS | 104.24 (42.93) | 25.85 (9.75) |
| REM | 61.29 (40.01) | 14.58 (8.70) |
| Sleep Parameter | Mean Minutes (SD) | Range |
| TST | 399.88 (68.06) | 203.5 – 472.5 |
| SOL | 15.24 (12.42) | 1 – 42.5 |
| WASO | 52.65 (56.47) | 0 - 225 |
| EEG Parameter | Mean (SD) |  |
| Mean Vector Length (MVL) | .23 (.01) | .21 - .27 |
| REM Theta Power | .20 (.04) | .02 - .33 |

**Note:** SD = standard deviation; TST = total sleep time; SOL = sleep onset latency; WASO = wake after sleep onset; N1 = stage 1; N2 = stage 2; SWS = slow wave sleep; REM = rapid eye movement sleep; MVL = PAC metric used to quantify the strength of coupling between SOs and spindles.

**Table 6.**
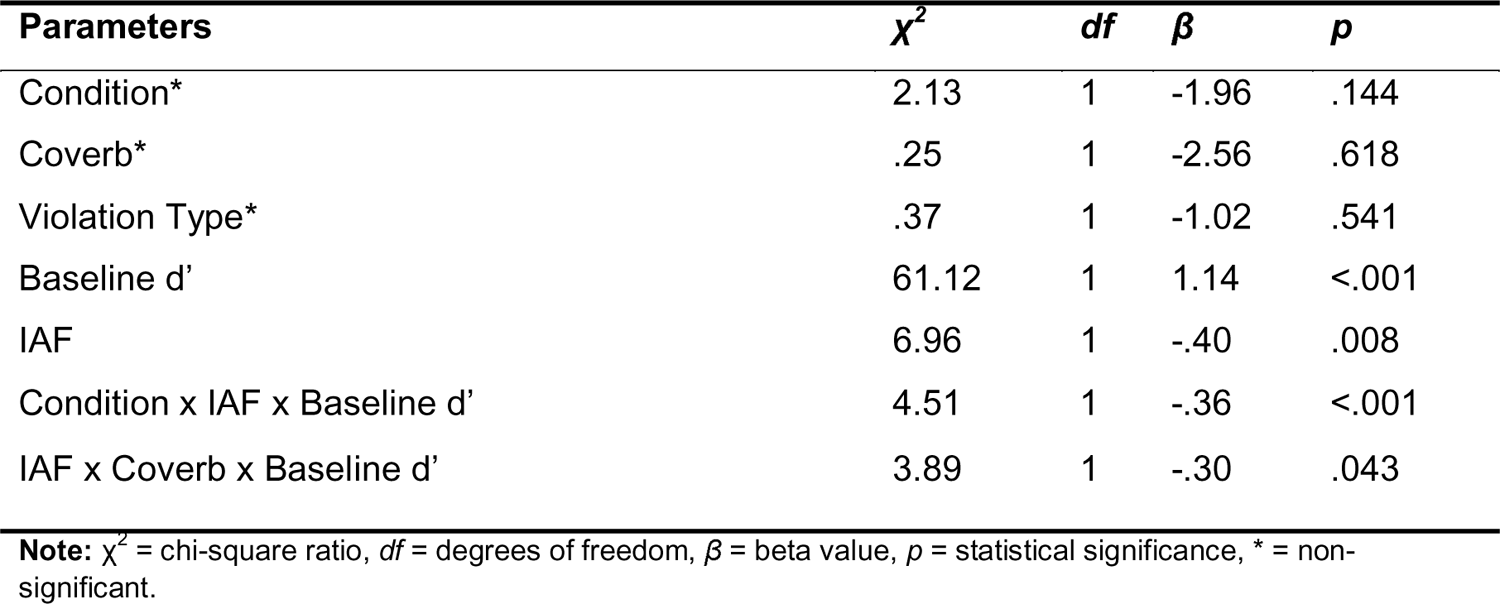
Statistical values for main effects and significant interaction effects within the behavioural model examined via type II Wald chi-square tests.

A significant interaction between IAF, Coverb and Baseline d’ was also revealed (statistical information can be found in Table 6 and the relationship is visualised in Appendix B [Figure 1A] of the supplementary material). Figure 4A illustrates that IAF differentially influenced memory performance at delayed testing depending both on condition and baseline performance. Further, sleep influenced memory performance, thus encouraging further examination of the effects of specific sleep physiology (i.e., PAC strength between SOs and spindles) on memory outcomes, which we report below.

### 3.3 Sleep Analysis

The results reported in the following summarise the effects most pertinent to the current hypotheses related to SWS neurophysiology, drawing on mixed model output summaries (for β values) and Type II Wald Chi-square tests for *p* values. Full model and Wald test summaries can be found in the supplementary material (Appendix A.2).

Mixed-effects modelling showed a significant interaction between PAC, Coverb and Violation Type, suggesting that coupling strength modulated performance at delayed testing, modulated by Coverb and Violation Type (refer to Table 7 for statistical values). As shown in Figure 4B, PAC did not influence performance for noun order violations of either coverb. However, for verb position violations, as coupling strength increased, differences in performance can be seen depending on coverb. Specifically, as coupling strength increased, performance for Undergoer-first constructions decreased and performance for Actor-first constructions increased.

**Table 7.** Statistical values for main effects and significant interaction effects within neurophysiological model examined via type II Wald chi-square tests.

| Parameters | $\chi^2$ | $df$ | $\beta$ | $p$ |
| --- | --- | --- | --- | --- |
| PAC* | .11 | 1 | .83 | .735 |
| IAF* | 1.76 | 1 | -.53 | .185 |
| Coverb* | 3.33 | 1 | -4.2 | .068 |
| Violation Type | 5.83 | 1 | -2.50 | .016 |
| Baseline $d'$ | 37.30 | 1 | -1.70 | <.001 |
| PAC x Coverb | 9.99 | 1 | -.30 | .001 |
| IAF x Coverb | 13.16 | 1 | .46 | <.001 |
| IAF x Violation Type | 6.71 | 1 | .25 | .009 |
| Coverb x Violation Type | 7.60 | 1 | 2.00 | .006 |
| IAF x Baseline $d'$ | 10.60 | 1 | .27 | .001 |
| PAC x Coverb x Violation Type | 5.91 | 1 | 1.30 | .015 |
| IAF x Coverb x Violation Type | 10.83 | 1 | -.23 | <.001 |
| IAF x Coverb x Baseline $d'$ | 9.42 | 1 | -.57 | .002 |
| IAF x Violation Type x Baseline $d'$ | 4.018 | 1 | -.28 | .045 |

**Note:** $\chi^2$ = chi-square ratio, $df$ = degrees of freedom, $\beta$ = beta value, $p$ = statistical significance, \* = non-significant.

Significant three-way interactions were also observed for (1) IAF x Coverb x Violation Type, (2) IAF x Coverb x Baseline d’, and (3) IAF x Violation Type x Baseline d’, indicating that IAF influences post-sleep memory performance, as modulated by grammatical rule type and pre-sleep performance (visualisations of these interactions can be found in Appendix B [Figure 1B, 1C and 1D, respectively] of the supplementary material. However, there was no signficiant interaction between PAC and IAF.

### 3.4 Exploratory Analysis

The results reported in the following summarise the effects most pertinent to exploratory investigation into REM sleep neurophysiology, drawing on mixed model output summaries (for β values) and Type II Wald Chi-square tests for *p* values. As literature suggests an effect of REM sleep on the consolidation of memory (Batterink et al., 2014; Djonlagic et al., 2009; Maquet et al., 2000), two exploratory models were run, with an initial model to investigate the effect of theta power during REM and a subsequent model exploring the effect of percentage of time spent in REM on memory. As the purpose of these models were to explore REM theta power and percentage of time spent in REM within a rule-based context, only interactions involving these predictors will be reported here. Full model and Wald test summaries for the REM theta power and percentage of REM models can be found in the supplementary material (Appendix A.3 & A.4, respectively).

The first mixed-model revealed a significant four-way REM Theta Power x IAF x Baseline d’ x Saggitality interaction, as shown in Figure 4C (refer to Table 8 for statistical values). Here, the largest differences between IAF values can be seen for higher pre-sleep performance scores and most prominently at posterior regions. There is also a notable relationship between post-sleep performance and theta power during REM for high IAF individuals, such that as theta power increases, memory performance also increases. The opposite pattern of results can be seen for low-IAF individuals. This pattern indicates both IAF and theta power modulate memory performance at delayed testing, and thus, rule extraction. Additionally, there was a significant REM Theta Power x IAF x Coverb x Saggitality interaction (statistical values can be found within Table 8 and this interaction is visualised within Appendix B [Figure 2A] of the supplementary material).

**Table 8.** Statistical values for main effects and significant interaction effects within the exploratory model (REM theta power) examined via type II Wald chi-square tests.

| Parameters | $\chi^2$ | df | $\beta$ | p |
| --- | --- | --- | --- | --- |
| REM Theta* | .00 | 1 | .54 | .997 |
| IAF* | 2.00 | 1 | -.67 | .157 |
| Saggitality* | .01 | 1 | .02, -.09 | .995 |
| Laterality* | .10 | 1 | -.10, .33 | .950 |
| Coverb | 9.84 | 1 | -2.40 | .002 |
| Violation Type | 21.93 | 1 | -1.80 | <.001 |
| Baseline d' | 185.85 | 1 | -4.30 | <.001 |
| REM Theta x Coverb | 8.30 | 1 | 3.25 | .004 |
| IAF x Coverb | 31.75 | 1 | .26 | <.001 |
| IAF x Violation Type | 33.87 | 1 | .16 | <.001 |
| Coverb x Violation Type | 53.21 | 1 | .32 | <.001 |
| IAF x Baseline d' | 110.88 | 1 | .55 | <.001 |
| Coverb x Baseline d' | 28.30 | 1 | 3.00 | <.001 |
| Violation Type x Baseline d' | 4.53 | 1 | 1.80 | .033 |
| IAF x Coverb x Violation Type | 18.72 | 1 | -.05 | <.001 |
| REM Theta x IAF x Baseline d' | 9.36 | 1 | .22 | .002 |
| IAF x Coverb x Baseline d' | 22.10 | 1 | -.36 | <.001 |
| IAF x Type x Baseline d' | 7.55 | 1 | -.17 | .006 |
| REM Theta x IAF x Coverb x Saggitality | 7.73 | 1 | .16, -.33 | .021 |
| REM Theta x IAF x Baseline d' x Saggitality | 6.66 | 1 | .04, -.33 | .036 |

**Note:** $\chi^2$ = chi-square ratio, df = degrees of freedom, $\beta$ = beta value, p = statistical significance, \* = non-significant.

The second mixed model explored percentage of time spent in REM sleep revealed a significant REM x Coverb x Type interaction (χ^2^ (1) = 7.41, β = −.69, p = .006), with the resolved interaction being shown in Figure 4D (refer to Table 9 for statistical values). Interestingly, percentage of time spent in REM also shows no visible effect on noun order violations. However, for verb positioning violations, there is a positive relationship between percentage of REM and post-sleep memory performance for Undergoer-first (bèi) constructions. In contrast, a negative relationship is seen for Actor-first (bǎ) constructions, indicating that processes during REM sleep are benefitting one rule type, whilst being detrimental for another. Additionally, there was a significant REM x Coverb x Baseline d’ interaction (statistical values can be found within Table 8 and this interaction is visualised within Appendix B [Figure 2B] of the supplementary material). However, there was no significant interaction between percentage of time spent in REM and IAF.

**Table 9.** Statistical values for main effects and significant interaction effects within the exploratory model (percentage of time in REM) examined via type II Wald chi-square tests.

| Parameters | $\chi^2$ | *df* | $\beta$ | *p* |
| --- | --- | --- | --- | --- |
| REM* | .02 | 1 | -4.20 | .888 |
| IAF* | 1.80 | 1 | -.27 | .180 |
| Coverb* | 2.85 | 1 | -7.2 | .091 |
| Violation Type | 4.43 | 1 | -1.20 | .035 |
| Baseline d' | 38.20 | 1 | 2.00 | <.001 |
| IAF x Coverb | 17.75 | 1 | .73 | <.001 |
| Coverb x Violation Type | 8.08 | 1 | 3.01 | .004 |
| IAF x Baseline d' | 5.45 | 1 | -0.06 | .020 |
| IAF x Coverb x Violation Type | 12.57 | 1 | -.32 | <.001 |
| REM x Coverb x Violation Type | 7.41 | 1 | -.69 | .006 |
| REM x Coverb x Baseline d' | 4.33 | 1 | 3.72 | .037 |
| IAF x Coverb x Baseline d' | 10.66 | 1 | -.86 | .001 |

**Note:** $\chi^2$ = chi-square ratio, *df* = degrees of freedom, $\beta$ = beta value, *p* = statistical significance, \* = non-significant.

## 4 DISCUSSION

Slow oscillation-spindle coupling during SWS and REM sleep have been independently linked to memory consolidation (Diekelmann & Born, 2010; Klinzing et al., 2019; Lewis & Durrant, 2011). Here, we provide evidence that these factors modulate memory performance within a rule-based context in language processing. Behaviourally, results indicate a positive relationship between baseline performance and delayed memory performance in both sleep and wake conditions that is modulated by individual alpha frequency (IAF). Regarding sleep neurophysiology, results indicate an interaction between phase-amplitude coupling (PAC) and sentence type (violation type and coverb). Specifically, higher PAC was associated with increased acceptability judgement performance for Actor-first (bǎ) sentences and decreased performance for Undergoer-first (bèi) with respect to the verb position rule. Our exploratory analyses of post-sleep memory performance further revealed a significant interaction between REM theta power, IAF, saggitality (anterior, midline and posterior regions) and pre-sleep performance. For high pre-sleep performers with a high IAF, as REM theta power increased, so did post-sleep memory performance, with this effect being most prominent in posterior regions. Additionally, proportion of REM sleep interacted with coverb and violation type, such that greater time spent in REM sleep was associated with decreased post-sleep performance on Actor-first constructions and increased performance for Undergoer-first constructions with respect to the verb position rule. Overall, results indicate that sleep neurophysiology and IAF influence consolidation and subsequent performance at both behavioural and neural levels and that patterns of memory performance differ across distinct rule types.

### 4.1 The influence of IAF and sleep on memory performance at the behavioural level

Our results partially supported the hypothesis (H^1^) that behavioural performance should be enhanced post-sleep relative to post-wake, however this relationship is modulated by IAF. Specifically, there was a generally positive relationship between baseline and delayed performance across conditions. However, for higher-IAF individuals in the wake condition, this relationship was weaker. As such, this result divergences from previous work showing that high IAF is associated with better performance on an array of tasks, including those assessing memory (Doppelmayr et al., 2005; Klimesch, 1997, 1999). More recent results (e.g., Ociepka et al., 2022) indicate that the association between high IAF and greater performance on a variety of tasks is not as common as once believed. In fact, in another reanalysis of Cross et al., (2021) examining a different linguistic manipulation, Nalaye et al., (2022) found a negative relationship between IAF and performance on the Mini Pinyin grammaticality judgement task, with lower-IAF individuals performing better. In combination with our results, it appears that high IAF is not beneficial in every context.

The link between IAF and information processing (Ociepka et al., 2022; Samaha & Postle, 2015; Surwillo, 1963) may influence the process of sleep-dependent memory consolidation which relies on hippocampal-to-neocortical information transfer (Yassa & Reagh, 2013). High IAF individuals in the wake condition may have been more susceptible to interference (Radvansky & Copeland, 2006) due to faster cycles of processing associated with higher IAF. Thus, lower IAF (slower cycles) may have been beneficial for those in the wake condition (Howard et al., 2017), with low IAF individuals preserving the initially encoded traces with less interference than high IAF individuals in the wake group. Our proposed role of IAF in information processing may also help to explain why baseline performance is critical when observing effects of IAF. It may be that the temporal receptive windows associated with IAF influence both the process of consolidation as well as initial encoding (Khader et al., 2010; Klimesch, 1997; Klimesch et al., 1996). However, this analysis did not specifically look at the process of encoding which limits the understanding of what may be occurring during this time (for an in-depth analysis of the encoding phase, see Cross et al., 2022). Despite this, analyses reported here provide further evidence for an effect of IAF on memory consolidation, consistent with previous literature (Doppelmayr et al., 2005; Klimesch, 1997, 1999; Richard Clark et al., 2004).

### 4.2 Neural mechanisms underlying sleep-based consolidation of rules

Both the behavioural and sleep analyses provide support for H^2^(a) which hypothesised that both IAF and PAC would predict post-sleep memory performance and that this relationship would be additive. Both models indicated that IAF influenced delayed memory performance. Further, in the sleep model, results showed an interaction between PAC, coverb, and violation type, such that as PAC increased, post-sleep memory performance for Actor-first sentences increased with respect to the verb position rule, while it decreased for Undergoer-first constructions. Slow oscillation-spindle coupling is suggested to reflect hippocampal-neocortical information transfer during NREM sleep (Diekelmann & Born, 2010; Klinzing et al., 2019; Lewis & Durrant, 2011), and as such, increased PAC associated with post-sleep performance suggests NREM sleep-based consolidation processes have occurred for rule-based information. This result is broadly consistent with previous literature linking NREM sleep neurophysiology to memory consolidation (Diekelmann & Born, 2010; Lewis & Durrant, 2011). This result also builds upon previous research linking SWS duration with rule extraction for both linguistic rules and non-linguistic rules (e.g., statistical; Batterink et al., 2014; Djonlagic et al., 2009; Durrant et al., 2013; Lerner & Gluck, 2019; Nieuwenhuis et al., 2013), linking specific PAC mechanisms (e.g., SO-spindle coupling) to memory performance in a rule-based context. In contrast, H^2^(b) was not supported, as PAC and IAF did not interact, contrasting previous literature in which IAF was associated with neurophysiology during SWS (Cross et al., 2020a).

Overall, PAC showed the greatest influence relative to the verb position rule, suggesting the novelty of a rule may play a role. Interestingly, there was no interaction present for noun order violations with both coverbs showing similar moderate performance (i.e., above chance). This may also suggest that rules related to noun order more generally were understood to some degree. However, when introducing the additional rule related to verb position, the role for PAC begins to form. When exploring the verb position rule, SO-spindle PAC demonstrated a differential effect depending on the coverb. Specifically, increased PAC moderately improved memory performance for Actor-first (bǎ) sentences. This may be due to native English speakers being more familiar with sentences in which the Actor occurs before the Undergoer (MacWhinney et al., 1984), which is also the case for bǎ sentences. As this rule type is less novel than the others examined here, better regularity extraction may have been influenced by such familiarity, leading to an effect of PAC for Actor-first sentences. By contrast, PAC negatively affected performance for verb position violations in Undergoer-first (bèi) sentences. This may have been due to the non-schema conforming nature of these constructions, which differ from English both with respect to noun order and verb position, resulting in a higher level of novelty. According to the iOTA model of sleep-based memory consolidation, non-schema conforming information is often downscaled to preferentially encode information analogous to existing knowledge (Lewis et al., 2018; Lewis & Durrant, 2011). Additionally, theories of memory consolidation, such as Complementary Learning Systems theory (McClelland et al., 1995), suggest that information similar to that of previous knowledge tends to be encoded and consolidated easier (Tse et al., 2011) and quicker (Tse et al., 2007), due to associative schemas. As such, the associative schema here is likely the grammatical elements of English, allowing the similar information (Actor-first sentences) to be more easily consolidated. Thus, this may explain how NREM mechanisms in our findings had the greatest effect on information closest to existing knowledge (i.e., Actor-first constructions) and had the opposite effect for information dissimilar to existing knowledge (i.e., Undergoer-first constructions), consistent with the iOTA model (Lewis et al., 2018; Lewis & Durrant, 2011).

### 4.3 The role of REM sleep in memory consolidation and rule extraction

As an exploratory step, we investigated the effect of REM sleep neurophysiology on rule extraction. Our results indicate differences in post-sleep memory performance relative to REM theta activity in posterior regions, contrasting with existing literature linking REM theta with central regions (Fogel et al., 2007) and anterior regions (Simor et al., 2016). Posterior theta has been proposed as the human correlate of the ponto-geniculo-occipital (PGO) wave (Frauscher et al., 2018), which has been previously attributed to memory consolidation and neuronal plasticity in experimental animal research (Datta, 2000; Datta et al., 2004). If so, the posterior theta activity reported here may reflect changes in neuronal plasticity (Grosmark et al., 2012; Hobson & Friston, 2012) and the consolidation of rule-based information. Additionally, the relationship between REM theta and post-sleep memory outcomes appears to differ depending on IAF. Our results revealed a positive relationship between REM theta and post-sleep memory for higher-IAF and a negative relationship for lower-IAF individuals. Cross et al., (2020a) also found a complex relationship between IAF and theta power during REM sleep. Higher IAF may have allowed for increased reactivations of memory during sleep or allowed the brain to preferentially encode salient information (Cross et al., 2020a; Heib et al., 2015).

We also observed an interaction between REM sleep percentage, coverb and violation type. Interestingly, noun order violations here also showed no interaction but moderate similar performance for both coverbs. However, when introducing the additional complexity related to the verb position rule, a relationship opposite to that seen for PAC can be observed. Specifically, higher proportions of REM sleep positively affected performance for Undergoer-first constructions (bèi) and led to decreased performance for Actor-first constructions (bǎ), contrasting with the pattern seen within NREM sleep mechanisms. This result partially converges with the results of Matorina & Poppenk, (2019), in which higher proportions of REM sleep led to a decrease in performance on a statistical learning task. Matorina & Poppenk, (2019) proposed that REM sleep may in fact be working against processes of extracting regularities, as it is more useful for a process called schema reconstruction, which disbands and reintegrates current and existing information into abstract schemas (Landmann et al., 2014). Further evidence for this comes from the iOTA model, in which REM sleep is proposed as the ideal setting for forming novel and abstract representations. Thus, NREM is likely most beneficial for associative memory processes, while REM sleep may facilitate the construction of abstract schemas (Lewis et al., 2018). The findings presented here provide support for this hypothesis, as demonstrated by the favourable effect of NREM sleep on Actor-first constructions (lower novelty) and REM sleep for Undergoer-first constructions (higher novelty). Overall, despite tentative interpretation, our results provide preliminary evidence for REM theta and IAF in rule-based contexts and demonstrate support for REM in non-schema conforming memory consolidation, providing a foundation for future research.

### 4.4 Limitations

As previously discussed, there are several limitations that should be addressed in future work. Firstly, analysis of electrophysiological activity during the encoding phase as they relate to delayed memory outcomes would provide additional understanding of the mechanisms underlying rule-based learning and consolidation. Additionally, whilst the focus here was on SO-spindle coupling, future research would benefit from including a broader range of sleep physiology metrics such as spindle-ripple coupling; albeit, this would need to be performed with invasive monitoring techniques, such as intracranial EEG, as scalp-recorded EEG is not capable of capturing activity in the ripple range (i.e., ∼ 80 – 140 Hz). This would aid in further understanding of the specific mechanisms driving rule-based consolidation during sleep. Additionally, the differing times of the day at which participants were tested may warrant further exploration into differences in circadian rhythms and any potential influence on wake-EEG (Aeschbach et al., 1997, 1999) and IAF. A basic comparison of baseline performance between sleep and wake revealed no meaningful differences in behavioural performance (refer to Appendix D to view output), suggesting that differing testing times may not have influenced performance in either group. Nonetheless, the relationship between IAF and circadian rhythms more specifically is largely unexplored and warrants further research. Despite these limitations, the current study provides a novel exploration of the neurophysiological mechanisms underlying the consolidation of rule-based information.

### 4.5 Conclusions

Here, we demonstrated that individual differences in intrinsic neural activity, indexed by IAF, predict rule extraction and consolidation across sleep and wake. Contrary to previous understanding of individual electrophysiological differences, we found a beneficial influence of lower IAF on memory within a rule-based context, proposing that higher-IAF and lower-IAF individuals draw on different information processing strategies as opposed to simply showing differing performance outcomes. We also demonstrated differential effects of SO-spindle coupling for specific grammatical rules, favouring the consolidation of those analogous to existing grammatical knowledge. We further provide support for the role of REM sleep in consolidating non-schema conforming information, potentially through schema reconstruction and the proposed role for REM for forming novel and abstract representations. Finally, we provided preliminary evidence for a relationship between REM sleep and IAF within rule extraction, a previously limited area of research. Future research should focus on further delineating the relationship between REM theta activity and individual processing capabilities in rule-based consolidation, which would further our understanding of the complex relationship between individual differences, sleep neurophysiology and memory consolidation.

## Funding and Acknowledgements

This work was supported by the Research Network for Undersea Decision Superiority 2020 Honours Scholarship awarded to MR and by an Australian Research Council Future Fellowship (FT160100437) awarded to IBS. ZC was also supported by Australian Commonwealth Government funding under the Research Training Program (RTP; number 212190) and Maurice de Rohan International Scholarship awarded to ZC. The authors would like to thank the research assistants Lena Zou, Erica Wilkinson, Nicole Vass, and Angela Osborn for help with data collection.

