## Supplementary Material for "Individual differences in information processing during sleep and wake predict sleep-based memory consolidation of complex rules"

July 25, 2023

Appendix A.1. Model summary and Wald Test output for behavioural model

Table 1: Wald Test output for behavioural model

| | Chisq | Df | Pr(>Chisq) |
|---|---|---|---|
| Condition | 2.13 | 1 | 0.14 |
| b_dprime | 61.12 | 1 | 0.00 |
| coverb | 0.25 | 1 | 0.62 |
| type | 0.37 | 1 | 0.54 |
| iaf | 6.96 | 1 | 0.01 |
| Condition:b_dprime | 0.03 | 1 | 0.85 |
| Condition:coverb | 1.62 | 1 | 0.20 |
| b_dprime:coverb | 1.92 | 1 | 0.17 |
| Condition:type | 0.52 | 1 | 0.47 |
| b_dprime:type | 0.00 | 1 | 0.95 |
| coverb:type | 1.13 | 1 | 0.29 |
| Condition:iaf | 0.13 | 1 | 0.72 |
| b_dprime:iaf | 1.48 | 1 | 0.22 |
| coverb:iaf | 3.26 | 1 | 0.07 |
| type:iaf | 1.76 | 1 | 0.18 |
| Condition:b_dprime:coverb | 0.36 | 1 | 0.55 |
| Condition:b_dprime:type | 1.05 | 1 | 0.30 |
| Condition:coverb:type | 2.22 | 1 | 0.14 |
| b_dprime:coverb:type | 0.17 | 1 | 0.68 |
| Condition:b_dprime:iaf | 10.89 | 1 | 0.00 |
| Condition:coverb:iaf | 0.02 | 1 | 0.88 |
| b_dprime:coverb:iaf | 4.08 | 1 | 0.04 |
| Condition:type:iaf | 1.45 | 1 | 0.23 |
| b_dprime:type:iaf | 2.35 | 1 | 0.13 |
| coverb:type:iaf | 3.09 | 1 | 0.08 |
| Condition:b_dprime:coverb:type | 0.01 | 1 | 0.92 |
| Condition:b_dprime:coverb:iaf | 0.70 | 1 | 0.40 |
| Condition:b_dprime:type:iaf | 0.01 | 1 | 0.92 |
| Condition:coverb:type:iaf | 0.63 | 1 | 0.43 |
| b_dprime:coverb:type:iaf | 0.03 | 1 | 0.87 |
| Condition:b_dprime:coverb:type:iaf | 0.31 | 1 | 0.58 |

Table 2: Model output for behavioural model

| | Estimate | Std. Error | t value |
|---|---|---|---|
| (Intercept) | 3.86 | 1.72 | 2.24 |
| Condition[1] | -1.96 | 1.72 | -1.14 |
| b_dprime | 1.14 | 1.40 | 0.82 |
| coverb[ba] | -2.56 | 1.18 | -2.16 |
| type[noun] | -1.02 | 1.24 | -0.82 |
| iaf | -0.39 | 0.18 | -2.21 |
| Condition[1]:b_dprime | 3.28 | 1.40 | 2.34 |
| Condition[1]:coverb[ba] | 0.65 | 1.18 | 0.55 |
| b_dprime:coverb[ba] | 2.74 | 1.43 | 1.92 |
| Condition[1]:type[noun] | 1.26 | 1.24 | 1.02 |
| b_dprime:type[noun] | 1.79 | 1.27 | 1.41 |
| coverb[ba]:type[noun] | 1.03 | 1.15 | 0.90 |
| Condition[1]:iaf | 0.19 | 0.18 | 1.05 |
| b_dprime:iaf | -0.03 | 0.15 | -0.18 |
| coverb[ba]:iaf | 0.28 | 0.12 | 2.31 |
| type[noun]:iaf | 0.10 | 0.13 | 0.81 |
| Condition[1]:b_dprime:coverb[ba] | -1.37 | 1.43 | -0.96 |
| Condition[1]:b_dprime:type[noun] | -0.49 | 1.27 | -0.39 |
| Condition[1]:coverb[ba]:type[noun] | -0.38 | 1.15 | -0.33 |
| b_dprime:coverb[ba]:type[noun] | 0.31 | 1.28 | 0.25 |
| Condition[1]:b_dprime:iaf | -0.36 | 0.15 | -2.41 |
| Condition[1]:coverb[ba]:iaf | -0.08 | 0.12 | -0.65 |
| b_dprime:coverb[ba]:iaf | -0.30 | 0.15 | -1.96 |
| Condition[1]:type[noun]:iaf | -0.12 | 0.13 | -0.97 |
| b_dprime:type[noun]:iaf | -0.19 | 0.14 | -1.42 |
| coverb[ba]:type[noun]:iaf | -0.11 | 0.12 | -0.97 |
| Condition[1]:b_dprime:coverb[ba]:type[noun] | -0.69 | 1.28 | -0.54 |
| Condition[1]:b_dprime:coverb[ba]:iaf | 0.15 | 0.15 | 1.00 |
| Condition[1]:b_dprime:type[noun]:iaf | 0.04 | 0.14 | 0.30 |
| Condition[1]:coverb[ba]:type[noun]:iaf | 0.05 | 0.12 | 0.44 |
| b_dprime:coverb[ba]:type[noun]:iaf | -0.03 | 0.14 | -0.20 |
| Condition[1]:b_dprime:coverb[ba]:type[noun]:iaf | 0.07 | 0.14 | 0.55 |

Appendix A.2. Model summary and Wald Test output for PAC model

Table 3: Wald Test output for PAC model

| | Chisq | Df | Pr(>Chisq) |
|---|---|---|---|
| Condition | 2.13 | 1 | 0.14 |
| b_dprime | 61.12 | 1 | 0.00 |
| coverb | 0.25 | 1 | 0.62 |
| type | 0.37 | 1 | 0.54 |
| iaf | 6.96 | 1 | 0.01 |
| Condition:b_dprime | 0.03 | 1 | 0.85 |
| Condition:coverb | 1.62 | 1 | 0.20 |
| b_dprime:coverb | 1.92 | 1 | 0.17 |
| Condition:type | 0.52 | 1 | 0.47 |
| b_dprime:type | 0.00 | 1 | 0.95 |
| coverb:type | 1.13 | 1 | 0.29 |
| Condition:iaf | 0.13 | 1 | 0.72 |
| b_dprime:iaf | 1.48 | 1 | 0.22 |
| coverb:iaf | 3.26 | 1 | 0.07 |
| type:iaf | 1.76 | 1 | 0.18 |
| Condition:b_dprime:coverb | 0.36 | 1 | 0.55 |
| Condition:b_dprime:type | 1.05 | 1 | 0.30 |
| Condition:coverb:type | 2.22 | 1 | 0.14 |
| b_dprime:coverb:type | 0.17 | 1 | 0.68 |
| Condition:b_dprime:iaf | 10.89 | 1 | 0.00 |
| Condition:coverb:iaf | 0.02 | 1 | 0.88 |
| b_dprime:coverb:iaf | 4.08 | 1 | 0.04 |
| Condition:type:iaf | 1.45 | 1 | 0.23 |
| b_dprime:type:iaf | 2.35 | 1 | 0.13 |
| coverb:type:iaf | 3.09 | 1 | 0.08 |
| Condition:b_dprime:coverb:type | 0.01 | 1 | 0.92 |
| Condition:b_dprime:coverb:iaf | 0.70 | 1 | 0.40 |
| Condition:b_dprime:type:iaf | 0.01 | 1 | 0.92 |
| Condition:coverb:type:iaf | 0.63 | 1 | 0.43 |
| b_dprime:coverb:type:iaf | 0.03 | 1 | 0.87 |
| Condition:b_dprime:coverb:type:iaf | 0.31 | 1 | 0.58 |

Table 4: Model output for PAC model

| | Estimate | Std. Error | t value |
|---|---|---|---|
| (Intercept) | 5.46 | 2.27 | 2.40 |
| scale(ndPAC) | 0.83 | 1.16 | 0.72 |
| iaf | -0.53 | 0.23 | -2.32 |
| coverb[ba] | -4.18 | 0.93 | -4.47 |
| type[noun] | -2.51 | 0.86 | -2.90 |
| b_dprime | -1.69 | 1.47 | -1.15 |
| scale(ndPAC):iaf | -0.09 | 0.13 | -0.73 |
| scale(ndPAC):coverb[ba] | -0.30 | 0.75 | -0.40 |
| iaf:coverb[ba] | 0.46 | 0.10 | 4.78 |
| scale(ndPAC):type[noun] | -0.66 | 0.73 | -0.90 |
| iaf:type[noun] | 0.25 | 0.09 | 2.82 |
| coverb[ba]:type[noun] | 1.96 | 0.85 | 2.29 |
| scale(ndPAC):b_dprime | -1.84 | 1.61 | -1.15 |
| iaf:b_dprime | 0.27 | 0.15 | 1.78 |
| coverb[ba]:b_dprime | 5.28 | 1.87 | 2.82 |
| type[noun]:b_dprime | 2.73 | 1.38 | 1.98 |
| scale(ndPAC):iaf:coverb[ba] | 0.06 | 0.08 | 0.75 |
| scale(ndPAC):iaf:type[noun] | 0.09 | 0.08 | 1.12 |
| scale(ndPAC):coverb[ba]:type[noun] | 1.27 | 0.73 | 1.74 |
| iaf:coverb[ba]:type[noun] | -0.23 | 0.09 | -2.60 |
| scale(ndPAC):iaf:b_dprime | 0.18 | 0.17 | 1.09 |
| scale(ndPAC):coverb[ba]:b_dprime | 0.38 | 1.57 | 0.24 |
| iaf:coverb[ba]:b_dprime | -0.57 | 0.20 | -2.90 |
| scale(ndPAC):type[noun]:b_dprime | 1.49 | 1.44 | 1.03 |
| iaf:type[noun]:b_dprime | -0.28 | 0.14 | -1.96 |
| coverb[ba]:type[noun]:b_dprime | 0.69 | 1.50 | 0.46 |
| scale(ndPAC):iaf:coverb[ba]:type[noun] | -0.16 | 0.08 | -1.99 |
| scale(ndPAC):iaf:coverb[ba]:b_dprime | -0.05 | 0.17 | -0.30 |
| scale(ndPAC):iaf:type[noun]:b_dprime | -0.17 | 0.15 | -1.10 |
| scale(ndPAC):coverb[ba]:type[noun]:b_dprime | -1.22 | 1.61 | -0.75 |
| iaf:coverb[ba]:type[noun]:b_dprime | -0.07 | 0.16 | -0.47 |
| scale(ndPAC):iaf:coverb[ba]:type[noun]:b_dprime | 0.12 | 0.17 | 0.72 |

Appendix A.3. Model summary and Wald Test output for REM theta model

Table 5: Wald Test output for REM theta model

| | Chisq | Df | Pr(>Chisq) |
|---|---|---|---|
| scale(Theta) | 0.00 | 1 | 1.00 |
| iaf | 2.00 | 1 | 0.16 |
| coverb | 9.84 | 1 | 0.00 |
| type | 21.94 | 1 | 0.00 |
| b_dprime | 185.83 | 1 | 0.00 |
| sag | 0.01 | 2 | 1.00 |
| lat | 0.10 | 2 | 0.95 |
| scale(Theta):iaf | 0.01 | 1 | 0.94 |
| scale(Theta):coverb | 8.31 | 1 | 0.00 |
| iaf:coverb | 31.75 | 1 | 0.00 |
| scale(Theta):type | 1.53 | 1 | 0.22 |
| iaf:type | 33.88 | 1 | 0.00 |
| coverb:type | 53.21 | 1 | 0.00 |
| scale(Theta):b_dprime | 0.72 | 1 | 0.40 |
| iaf:b_dprime | 110.88 | 1 | 0.00 |
| coverb:b_dprime | 28.31 | 1 | 0.00 |
| type:b_dprime | 4.53 | 1 | 0.03 |
| scale(Theta):sag | 0.42 | 2 | 0.81 |
| iaf:sag | 0.18 | 2 | 0.91 |
| coverb:sag | 0.06 | 2 | 0.97 |
| type:sag | 0.42 | 2 | 0.81 |
| b_dprime:sag | 0.08 | 2 | 0.96 |
| scale(Theta):lat | 0.02 | 2 | 0.99 |
| iaf:lat | 0.01 | 2 | 1.00 |
| coverb:lat | 0.06 | 2 | 0.97 |
| type:lat | 0.10 | 2 | 0.95 |
| b_dprime:lat | 0.03 | 2 | 0.99 |
| sag:lat | 0.04 | 4 | 1.00 |
| scale(Theta):iaf:coverb | 0.02 | 1 | 0.89 |
| scale(Theta):iaf:type | 1.90 | 1 | 0.17 |
| scale(Theta):coverb:type | 0.30 | 1 | 0.58 |
| iaf:coverb:type | 18.72 | 1 | 0.00 |
| scale(Theta):iaf:b_dprime | 9.36 | 1 | 0.00 |
| scale(Theta):coverb:b_dprime | 0.67 | 1 | 0.41 |
| iaf:coverb:b_dprime | 22.10 | 1 | 0.00 |
| scale(Theta):type:b_dprime | 0.39 | 1 | 0.53 |
| iaf:type:b_dprime | 7.55 | 1 | 0.01 |
| coverb:type:b_dprime | 0.05 | 1 | 0.82 |
| scale(Theta):iaf:sag | 0.01 | 2 | 1.00 |
| scale(Theta):coverb:sag | 3.02 | 2 | 0.22 |
| iaf:coverb:sag | 0.76 | 2 | 0.69 |
| scale(Theta):type:sag | 0.38 | 2 | 0.83 |
| iaf:type:sag | 1.64 | 2 | 0.44 |
| coverb:type:sag | 0.41 | 2 | 0.81 |
| scale(Theta):b_dprime:sag | 1.99 | 2 | 0.37 |
| iaf:b_dprime:sag | 0.93 | 2 | 0.63 |
| coverb:b_dprime:sag | 0.90 | 2 | 0.64 |
| type:b_dprime:sag | 0.23 | 2 | 0.89 |
| scale(Theta):iaf:lat | 0.03 | 2 | 0.98 |
| scale(Theta):coverb:lat | 0.86 | 2 | 0.65 |
| iaf:coverb:lat | 0.64 | 2 | 0.73 |
| scale(Theta):type:lat | 1.79 | 2 | 0.41 |
| iaf:type:lat | 1.95 | 2 | 0.38 |
| coverb:type:lat | 0.49 | 2 | 0.78 |
| scale(Theta):b_dprime:lat | 0.13 | 2 | 0.94 |

Table 5: Wald Test output for REM theta model *(continued)*

| | Chisq | Df | Pr(>Chisq) |
|---|---|---|---|
| iaf:b_dprime:lat | 0.15 | 2 | 0.93 |
| coverb:b_dprime:lat | 1.46 | 2 | 0.48 |
| type:b_dprime:lat | 0.63 | 2 | 0.73 |
| scale(Theta):sag:lat | 0.52 | 4 | 0.97 |
| iaf:sag:lat | 0.27 | 4 | 0.99 |
| coverb:sag:lat | 0.08 | 4 | 1.00 |
| type:sag:lat | 0.28 | 4 | 0.99 |
| b_dprime:sag:lat | 0.69 | 4 | 0.95 |
| scale(Theta):iaf:coverb:type | 0.29 | 1 | 0.59 |
| scale(Theta):iaf:coverb:b_dprime | 0.36 | 1 | 0.55 |
| scale(Theta):iaf:type:b_dprime | 0.37 | 1 | 0.54 |
| scale(Theta):coverb:type:b_dprime | 0.53 | 1 | 0.47 |
| iaf:coverb:type:b_dprime | 2.83 | 1 | 0.09 |
| scale(Theta):iaf:coverb:sag | 7.73 | 2 | 0.02 |
| scale(Theta):iaf:type:sag | 3.84 | 2 | 0.15 |
| scale(Theta):coverb:type:sag | 4.59 | 2 | 0.10 |
| iaf:coverb:type:sag | 1.42 | 2 | 0.49 |
| scale(Theta):iaf:b_dprime:sag | 6.67 | 2 | 0.04 |
| scale(Theta):coverb:b_dprime:sag | 3.31 | 2 | 0.19 |
| iaf:coverb:b_dprime:sag | 0.80 | 2 | 0.67 |
| scale(Theta):type:b_dprime:sag | 0.51 | 2 | 0.78 |
| iaf:type:b_dprime:sag | 0.37 | 2 | 0.83 |
| coverb:type:b_dprime:sag | 1.02 | 2 | 0.60 |
| scale(Theta):iaf:coverb:lat | 1.77 | 2 | 0.41 |
| scale(Theta):iaf:type:lat | 5.02 | 2 | 0.08 |
| scale(Theta):coverb:type:lat | 0.30 | 2 | 0.86 |
| iaf:coverb:type:lat | 0.74 | 2 | 0.69 |
| scale(Theta):iaf:b_dprime:lat | 0.49 | 2 | 0.78 |
| scale(Theta):coverb:b_dprime:lat | 0.86 | 2 | 0.65 |
| iaf:coverb:b_dprime:lat | 0.36 | 2 | 0.83 |
| scale(Theta):type:b_dprime:lat | 1.64 | 2 | 0.44 |
| iaf:type:b_dprime:lat | 1.92 | 2 | 0.38 |
| coverb:type:b_dprime:lat | 0.04 | 2 | 0.98 |
| scale(Theta):iaf:sag:lat | 0.30 | 4 | 0.99 |
| scale(Theta):coverb:sag:lat | 1.45 | 4 | 0.84 |
| iaf:coverb:sag:lat | 2.52 | 4 | 0.64 |
| scale(Theta):type:sag:lat | 2.10 | 4 | 0.72 |
| iaf:type:sag:lat | 2.47 | 4 | 0.65 |
| coverb:type:sag:lat | 0.99 | 4 | 0.91 |
| scale(Theta):b_dprime:sag:lat | 0.32 | 4 | 0.99 |
| iaf:b_dprime:sag:lat | 0.22 | 4 | 0.99 |
| coverb:b_dprime:sag:lat | 1.08 | 4 | 0.90 |
| type:b_dprime:sag:lat | 1.22 | 4 | 0.87 |
| scale(Theta):iaf:coverb:type:b_dprime | 1.73 | 1 | 0.19 |
| scale(Theta):iaf:coverb:type:sag | 4.38 | 2 | 0.11 |
| scale(Theta):iaf:coverb:b_dprime:sag | 1.46 | 2 | 0.48 |
| scale(Theta):iaf:type:b_dprime:sag | 1.92 | 2 | 0.38 |
| scale(Theta):coverb:type:b_dprime:sag | 2.12 | 2 | 0.35 |
| iaf:coverb:type:b_dprime:sag | 0.49 | 2 | 0.78 |
| scale(Theta):iaf:coverb:type:lat | 5.02 | 2 | 0.08 |
| scale(Theta):iaf:coverb:b_dprime:lat | 3.06 | 2 | 0.22 |
| scale(Theta):iaf:type:b_dprime:lat | 3.90 | 2 | 0.14 |
| scale(Theta):coverb:type:b_dprime:lat | 2.56 | 2 | 0.28 |
| iaf:coverb:type:b_dprime:lat | 0.13 | 2 | 0.94 |
| scale(Theta):iaf:coverb:sag:lat | 0.34 | 4 | 0.99 |
| scale(Theta):iaf:type:sag:lat | 1.34 | 4 | 0.85 |
| scale(Theta):coverb:type:sag:lat | 0.80 | 4 | 0.94 |

Table 5: Wald Test output for REM theta model *(continued)*

| | Chisq | Df | Pr(>Chisq) |
|---|---|---|---|
| iaf:coverb:type:sag:lat | 1.25 | 4 | 0.87 |
| scale(Theta):iaf:b_dprime:sag:lat | 0.53 | 4 | 0.97 |
| scale(Theta):coverb:b_dprime:sag:lat | 0.80 | 4 | 0.94 |
| iaf:coverb:b_dprime:sag:lat | 0.62 | 4 | 0.96 |
| scale(Theta):type:b_dprime:sag:lat | 1.14 | 4 | 0.89 |
| iaf:type:b_dprime:sag:lat | 0.87 | 4 | 0.93 |
| coverb:type:b_dprime:sag:lat | 0.72 | 4 | 0.95 |
| scale(Theta):iaf:coverb:type:b_dprime:sag | 1.46 | 2 | 0.48 |
| scale(Theta):iaf:coverb:type:b_dprime:lat | 4.03 | 2 | 0.13 |
| scale(Theta):iaf:coverb:type:sag:lat | 0.57 | 4 | 0.97 |
| scale(Theta):iaf:coverb:b_dprime:sag:lat | 0.78 | 4 | 0.94 |
| scale(Theta):iaf:type:b_dprime:sag:lat | 1.30 | 4 | 0.86 |
| scale(Theta):coverb:type:b_dprime:sag:lat | 1.21 | 4 | 0.88 |
| iaf:coverb:type:b_dprime:sag:lat | 1.12 | 4 | 0.89 |
| scale(Theta):iaf:coverb:type:b_dprime:sag:lat | 1.30 | 4 | 0.86 |

Table 6: Model output for REM theta model

| | Estimate | Std. Error | t value |
|---|---|---|---|
| (Intercept) | 6.75 | 2.21 | 3.05 |
| scale(Theta) | 0.54 | 1.12 | 0.48 |
| iaf | -0.67 | 0.23 | -2.97 |
| coverb[ba] | -2.37 | 0.59 | -4.01 |
| type[noun] | -1.80 | 0.59 | -3.03 |
| b_dprime | -4.31 | 1.03 | -4.18 |
| sag[anterior] | 0.03 | 0.82 | 0.03 |
| sag[central] | -0.09 | 0.82 | -0.10 |
| lat[left] | -0.10 | 0.88 | -0.11 |
| lat[midline] | 0.33 | 0.77 | 0.43 |
| scale(Theta):iaf | -0.06 | 0.12 | -0.50 |
| scale(Theta):coverb[ba] | 3.25 | 0.97 | 3.34 |
| iaf:coverb[ba] | 0.26 | 0.06 | 4.11 |
| scale(Theta):type[noun] | 3.02 | 0.93 | 3.23 |
| iaf:type[noun] | 0.16 | 0.06 | 2.47 |
| coverb[ba]:type[noun] | 0.32 | 0.57 | 0.55 |
| scale(Theta):b_dprime | -2.06 | 1.48 | -1.39 |
| iaf:b_dprime | 0.55 | 0.11 | 5.15 |
| coverb[ba]:b_dprime | 3.01 | 1.12 | 2.69 |
| type[noun]:b_dprime | 1.77 | 1.03 | 1.72 |
| scale(Theta):sag[anterior] | 0.68 | 1.30 | 0.52 |
| scale(Theta):sag[central] | -1.30 | 1.33 | -0.98 |
| iaf:sag[anterior] | -0.01 | 0.09 | -0.08 |
| iaf:sag[central] | 0.01 | 0.09 | 0.17 |
| coverb[ba]:sag[anterior] | 0.27 | 0.82 | 0.33 |
| coverb[ba]:sag[central] | 0.41 | 0.80 | 0.51 |
| type[noun]:sag[anterior] | 0.56 | 0.82 | 0.68 |
| type[noun]:sag[central] | -0.99 | 0.80 | -1.24 |
| b_dprime:sag[anterior] | 0.03 | 1.31 | 0.02 |
| b_dprime:sag[central] | -0.24 | 1.53 | -0.16 |
| scale(Theta):lat[left] | -1.05 | 1.39 | -0.75 |
| scale(Theta):lat[midline] | 1.50 | 1.17 | 1.28 |
| iaf:lat[left] | 0.01 | 0.10 | 0.14 |
| iaf:lat[midline] | -0.04 | 0.08 | -0.46 |
| coverb[ba]:lat[left] | 1.07 | 0.86 | 1.25 |
| coverb[ba]:lat[midline] | -0.70 | 0.75 | -0.94 |
| type[noun]:lat[left] | 1.66 | 0.85 | 1.95 |
| type[noun]:lat[midline] | -1.47 | 0.74 | -1.97 |
| b_dprime:lat[left] | 0.27 | 1.31 | 0.21 |
| b_dprime:lat[midline] | -0.78 | 1.42 | -0.55 |
| sag[anterior]:lat[left] | -0.08 | 1.29 | -0.06 |
| sag[central]:lat[left] | 0.03 | 1.19 | 0.03 |
| sag[anterior]:lat[midline] | -0.03 | 1.09 | -0.02 |
| sag[central]:lat[midline] | 0.25 | 1.14 | 0.22 |
| scale(Theta):iaf:coverb[ba] | -0.36 | 0.10 | -3.40 |
| scale(Theta):iaf:type[noun] | -0.33 | 0.10 | -3.26 |
| scale(Theta):coverb[ba]:type[noun] | -2.96 | 0.97 | -3.05 |
| iaf:coverb[ba]:type[noun] | -0.04 | 0.06 | -0.71 |
| scale(Theta):iaf:b_dprime | 0.22 | 0.16 | 1.43 |
| scale(Theta):coverb[ba]:b_dprime | -2.58 | 1.73 | -1.49 |
| iaf:coverb[ba]:b_dprime | -0.34 | 0.12 | -2.92 |
| scale(Theta):type[noun]:b_dprime | -1.55 | 1.40 | -1.10 |
| iaf:type[noun]:b_dprime | -0.17 | 0.11 | -1.64 |
| coverb[ba]:type[noun]:b_dprime | 1.17 | 1.07 | 1.09 |
| scale(Theta):iaf:sag[anterior] | -0.07 | 0.14 | -0.51 |
| scale(Theta):iaf:sag[central] | 0.14 | 0.14 | 1.00 |
| scale(Theta):coverb[ba]:sag[anterior] | -1.54 | 1.27 | -1.21 |

Table 6: Model output for REM theta model *(continued)*

| | Estimate | Std. Error | t value |
|---|---|---|---|
| scale(Theta):coverb[ba]:sag[central] | 2.98 | 1.28 | 2.34 |
| iaf:coverb[ba]:sag[anterior] | -0.03 | 0.09 | -0.29 |
| iaf:coverb[ba]:sag[central] | -0.05 | 0.08 | -0.59 |
| scale(Theta):type[noun]:sag[anterior] | -1.70 | 1.25 | -1.36 |
| scale(Theta):type[noun]:sag[central] | 2.06 | 1.28 | 1.61 |
| iaf:type[noun]:sag[anterior] | -0.05 | 0.09 | -0.55 |
| iaf:type[noun]:sag[central] | 0.09 | 0.08 | 1.13 |
| coverb[ba]:type[noun]:sag[anterior] | -0.22 | 0.81 | -0.27 |
| coverb[ba]:type[noun]:sag[central] | -0.14 | 0.80 | -0.17 |
| scale(Theta):b_dprime:sag[anterior] | -0.42 | 1.80 | -0.23 |
| scale(Theta):b_dprime:sag[central] | 3.09 | 1.84 | 1.68 |
| iaf:b_dprime:sag[anterior] | 0.00 | 0.14 | -0.01 |
| iaf:b_dprime:sag[central] | 0.02 | 0.15 | 0.10 |
| coverb[ba]:b_dprime:sag[anterior] | 0.73 | 1.32 | 0.55 |
| coverb[ba]:b_dprime:sag[central] | -1.02 | 1.54 | -0.67 |
| type[noun]:b_dprime:sag[anterior] | 0.04 | 1.30 | 0.03 |
| type[noun]:b_dprime:sag[central] | 0.40 | 1.49 | 0.27 |
| scale(Theta):iaf:lat[left] | 0.11 | 0.15 | 0.76 |
| scale(Theta):iaf:lat[midline] | -0.17 | 0.13 | -1.31 |
| scale(Theta):coverb[ba]:lat[left] | 2.29 | 1.38 | 1.66 |
| scale(Theta):coverb[ba]:lat[midline] | -2.34 | 1.13 | -2.08 |
| iaf:coverb[ba]:lat[left] | -0.13 | 0.09 | -1.39 |
| iaf:coverb[ba]:lat[midline] | 0.09 | 0.08 | 1.14 |
| scale(Theta):type[noun]:lat[left] | 3.07 | 1.37 | 2.23 |
| scale(Theta):type[noun]:lat[midline] | -3.20 | 1.12 | -2.86 |
| iaf:type[noun]:lat[left] | -0.19 | 0.09 | -2.05 |
| iaf:type[noun]:lat[midline] | 0.16 | 0.08 | 2.10 |
| coverb[ba]:type[noun]:lat[left] | -1.31 | 0.86 | -1.53 |
| coverb[ba]:type[noun]:lat[midline] | 1.04 | 0.74 | 1.40 |
| scale(Theta):b_dprime:lat[left] | 0.96 | 2.09 | 0.46 |
| scale(Theta):b_dprime:lat[midline] | -0.79 | 1.66 | -0.47 |
| iaf:b_dprime:lat[left] | -0.02 | 0.14 | -0.18 |
| iaf:b_dprime:lat[midline] | 0.08 | 0.14 | 0.54 |
| coverb[ba]:b_dprime:lat[left] | -0.56 | 1.31 | -0.43 |
| coverb[ba]:b_dprime:lat[midline] | -0.01 | 1.43 | -0.01 |
| type[noun]:b_dprime:lat[left] | -1.47 | 1.30 | -1.13 |
| type[noun]:b_dprime:lat[midline] | 1.36 | 1.42 | 0.96 |
| scale(Theta):sag[anterior]:lat[left] | -0.71 | 1.84 | -0.39 |
| scale(Theta):sag[central]:lat[left] | 0.99 | 1.89 | 0.52 |
| scale(Theta):sag[anterior]:lat[midline] | 0.14 | 1.64 | 0.09 |
| scale(Theta):sag[central]:lat[midline] | -0.19 | 1.61 | -0.12 |
| iaf:sag[anterior]:lat[left] | 0.01 | 0.14 | 0.08 |
| iaf:sag[central]:lat[left] | -0.01 | 0.13 | -0.06 |
| iaf:sag[anterior]:lat[midline] | 0.00 | 0.12 | -0.02 |
| iaf:sag[central]:lat[midline] | -0.02 | 0.12 | -0.19 |
| coverb[ba]:sag[anterior]:lat[left] | 1.01 | 1.29 | 0.79 |
| coverb[ba]:sag[central]:lat[left] | -0.15 | 1.19 | -0.13 |
| coverb[ba]:sag[anterior]:lat[midline] | -0.08 | 1.09 | -0.08 |
| coverb[ba]:sag[central]:lat[midline] | -0.16 | 1.14 | -0.14 |
| type[noun]:sag[anterior]:lat[left] | 0.89 | 1.29 | 0.69 |
| type[noun]:sag[central]:lat[left] | -0.12 | 1.19 | -0.10 |
| type[noun]:sag[anterior]:lat[midline] | 0.63 | 1.08 | 0.58 |
| type[noun]:sag[central]:lat[midline] | -0.65 | 1.14 | -0.57 |
| b_dprime:sag[anterior]:lat[left] | -0.57 | 1.82 | -0.31 |
| b_dprime:sag[central]:lat[left] | -0.07 | 1.96 | -0.04 |
| b_dprime:sag[anterior]:lat[midline] | 0.25 | 1.94 | 0.13 |
| b_dprime:sag[central]:lat[midline] | -0.29 | 2.35 | -0.12 |

Table 6: Model output for REM theta model *(continued)*

| | Estimate | Std. Error | t value |
|---|---|---|---|
| scale(Theta):iaf:coverb[ba]:type[noun] | 0.33 | 0.10 | 3.15 |
| scale(Theta):iaf:coverb[ba]:b_dprime | 0.28 | 0.18 | 1.56 |
| scale(Theta):iaf:type[noun]:b_dprime | 0.17 | 0.15 | 1.15 |
| scale(Theta):coverb[ba]:type[noun]:b_dprime | 0.71 | 1.41 | 0.50 |
| iaf:coverb[ba]:type[noun]:b_dprime | -0.12 | 0.11 | -1.12 |
| scale(Theta):iaf:coverb[ba]:sag[anterior] | 0.16 | 0.14 | 1.18 |
| scale(Theta):iaf:coverb[ba]:sag[central] | -0.33 | 0.14 | -2.43 |
| scale(Theta):iaf:type[noun]:sag[anterior] | 0.18 | 0.13 | 1.30 |
| scale(Theta):iaf:type[noun]:sag[central] | -0.21 | 0.14 | -1.56 |
| scale(Theta):coverb[ba]:type[noun]:sag[anterior] | 1.90 | 1.25 | 1.52 |
| scale(Theta):coverb[ba]:type[noun]:sag[central] | -2.22 | 1.28 | -1.74 |
| iaf:coverb[ba]:type[noun]:sag[anterior] | 0.02 | 0.09 | 0.25 |
| iaf:coverb[ba]:type[noun]:sag[central] | 0.02 | 0.08 | 0.21 |
| scale(Theta):iaf:b_dprime:sag[anterior] | 0.04 | 0.19 | 0.23 |
| scale(Theta):iaf:b_dprime:sag[central] | -0.33 | 0.19 | -1.71 |
| scale(Theta):coverb[ba]:b_dprime:sag[anterior] | 1.57 | 1.89 | 0.83 |
| scale(Theta):coverb[ba]:b_dprime:sag[central] | -0.82 | 1.86 | -0.44 |
| iaf:coverb[ba]:b_dprime:sag[anterior] | -0.08 | 0.14 | -0.55 |
| iaf:coverb[ba]:b_dprime:sag[central] | 0.11 | 0.16 | 0.68 |
| scale(Theta):type[noun]:b_dprime:sag[anterior] | 2.24 | 1.80 | 1.25 |
| scale(Theta):type[noun]:b_dprime:sag[central] | -1.09 | 1.84 | -0.60 |
| iaf:type[noun]:b_dprime:sag[anterior] | -0.01 | 0.13 | -0.09 |
| iaf:type[noun]:b_dprime:sag[central] | -0.04 | 0.15 | -0.27 |
| coverb[ba]:type[noun]:b_dprime:sag[anterior] | -0.76 | 1.31 | -0.58 |
| coverb[ba]:type[noun]:b_dprime:sag[central] | 0.51 | 1.50 | 0.34 |
| scale(Theta):iaf:coverb[ba]:lat[left] | -0.26 | 0.15 | -1.73 |
| scale(Theta):iaf:coverb[ba]:lat[midline] | 0.26 | 0.12 | 2.09 |
| scale(Theta):iaf:type[noun]:lat[left] | -0.35 | 0.15 | -2.33 |
| scale(Theta):iaf:type[noun]:lat[midline] | 0.36 | 0.12 | 2.94 |
| scale(Theta):coverb[ba]:type[noun]:lat[left] | -2.18 | 1.38 | -1.58 |
| scale(Theta):coverb[ba]:type[noun]:lat[midline] | 3.08 | 1.13 | 2.71 |
| iaf:coverb[ba]:type[noun]:lat[left] | 0.15 | 0.09 | 1.64 |
| iaf:coverb[ba]:type[noun]:lat[midline] | -0.13 | 0.08 | -1.65 |
| scale(Theta):iaf:b_dprime:lat[left] | -0.10 | 0.22 | -0.47 |
| scale(Theta):iaf:b_dprime:lat[midline] | 0.09 | 0.17 | 0.50 |
| scale(Theta):coverb[ba]:b_dprime:lat[left] | -1.37 | 2.11 | -0.65 |
| scale(Theta):coverb[ba]:b_dprime:lat[midline] | 1.83 | 1.71 | 1.07 |
| iaf:coverb[ba]:b_dprime:lat[left] | 0.07 | 0.14 | 0.52 |
| iaf:coverb[ba]:b_dprime:lat[midline] | -0.01 | 0.14 | -0.07 |
| scale(Theta):type[noun]:b_dprime:lat[left] | -2.16 | 2.09 | -1.04 |
| scale(Theta):type[noun]:b_dprime:lat[midline] | 2.72 | 1.66 | 1.64 |
| iaf:type[noun]:b_dprime:lat[left] | 0.16 | 0.14 | 1.22 |
| iaf:type[noun]:b_dprime:lat[midline] | -0.15 | 0.14 | -1.05 |
| coverb[ba]:type[noun]:b_dprime:lat[left] | 0.55 | 1.30 | 0.42 |
| coverb[ba]:type[noun]:b_dprime:lat[midline] | -0.39 | 1.41 | -0.27 |
| scale(Theta):iaf:sag[anterior]:lat[left] | 0.08 | 0.20 | 0.38 |
| scale(Theta):iaf:sag[central]:lat[left] | -0.11 | 0.20 | -0.52 |
| scale(Theta):iaf:sag[anterior]:lat[midline] | -0.01 | 0.18 | -0.08 |
| scale(Theta):iaf:sag[central]:lat[midline] | 0.02 | 0.17 | 0.09 |
| scale(Theta):coverb[ba]:sag[anterior]:lat[left] | 0.32 | 1.84 | 0.17 |
| scale(Theta):coverb[ba]:sag[central]:lat[left] | -0.65 | 1.89 | -0.34 |
| scale(Theta):coverb[ba]:sag[anterior]:lat[midline] | 1.24 | 1.62 | 0.77 |
| scale(Theta):coverb[ba]:sag[central]:lat[midline] | -0.41 | 1.61 | -0.25 |
| iaf:coverb[ba]:sag[anterior]:lat[left] | -0.11 | 0.14 | -0.80 |
| iaf:coverb[ba]:sag[central]:lat[left] | 0.02 | 0.13 | 0.14 |
| iaf:coverb[ba]:sag[anterior]:lat[midline] | 0.00 | 0.12 | 0.02 |
| iaf:coverb[ba]:sag[central]:lat[midline] | 0.03 | 0.12 | 0.22 |

Table 6: Model output for REM theta model *(continued)*

| | Estimate | Std. Error | t value |
|---|---|---|---|
| scale(Theta):type[noun]:sag[anterior]:lat[left] | 0.76 | 1.84 | 0.41 |
| scale(Theta):type[noun]:sag[central]:lat[left] | -1.36 | 1.89 | -0.72 |
| scale(Theta):type[noun]:sag[anterior]:lat[midline] | 1.58 | 1.62 | 0.97 |
| scale(Theta):type[noun]:sag[central]:lat[midline] | -0.38 | 1.61 | -0.24 |
| iaf:type[noun]:sag[anterior]:lat[left] | -0.11 | 0.14 | -0.75 |
| iaf:type[noun]:sag[central]:lat[left] | 0.02 | 0.13 | 0.19 |
| iaf:type[noun]:sag[anterior]:lat[midline] | -0.07 | 0.11 | -0.58 |
| iaf:type[noun]:sag[central]:lat[midline] | 0.06 | 0.12 | 0.56 |
| coverb[ba]:type[noun]:sag[anterior]:lat[left] | -0.83 | 1.29 | -0.65 |
| coverb[ba]:type[noun]:sag[central]:lat[left] | 0.62 | 1.19 | 0.52 |
| coverb[ba]:type[noun]:sag[anterior]:lat[midline] | 0.40 | 1.08 | 0.37 |
| coverb[ba]:type[noun]:sag[central]:lat[midline] | -0.49 | 1.14 | -0.43 |
| scale(Theta):b_dprime:sag[anterior]:lat[left] | 1.17 | 2.68 | 0.43 |
| scale(Theta):b_dprime:sag[central]:lat[left] | -1.42 | 2.73 | -0.52 |
| scale(Theta):b_dprime:sag[anterior]:lat[midline] | -0.71 | 2.30 | -0.31 |
| scale(Theta):b_dprime:sag[central]:lat[midline] | 0.99 | 2.33 | 0.43 |
| iaf:b_dprime:sag[anterior]:lat[left] | 0.06 | 0.19 | 0.29 |
| iaf:b_dprime:sag[central]:lat[left] | 0.01 | 0.20 | 0.05 |
| iaf:b_dprime:sag[anterior]:lat[midline] | -0.01 | 0.20 | -0.07 |
| iaf:b_dprime:sag[central]:lat[midline] | 0.02 | 0.23 | 0.09 |
| coverb[ba]:b_dprime:sag[anterior]:lat[left] | -1.00 | 1.82 | -0.55 |
| coverb[ba]:b_dprime:sag[central]:lat[left] | 0.56 | 1.97 | 0.28 |
| coverb[ba]:b_dprime:sag[anterior]:lat[midline] | 0.03 | 1.94 | 0.01 |
| coverb[ba]:b_dprime:sag[central]:lat[midline] | -0.92 | 2.38 | -0.39 |
| type[noun]:b_dprime:sag[anterior]:lat[left] | -0.44 | 1.82 | -0.24 |
| type[noun]:b_dprime:sag[central]:lat[left] | -0.29 | 1.96 | -0.15 |
| type[noun]:b_dprime:sag[anterior]:lat[midline] | -1.43 | 1.93 | -0.74 |
| type[noun]:b_dprime:sag[central]:lat[midline] | 1.04 | 2.35 | 0.45 |
| scale(Theta):iaf:coverb[ba]:type[noun]:b_dprime | -0.08 | 0.15 | -0.57 |
| scale(Theta):iaf:coverb[ba]:type[noun]:sag[anterior] | -0.20 | 0.14 | -1.44 |
| scale(Theta):iaf:coverb[ba]:type[noun]:sag[central] | 0.25 | 0.14 | 1.83 |
| scale(Theta):iaf:coverb[ba]:b_dprime:sag[anterior] | -0.17 | 0.20 | -0.85 |
| scale(Theta):iaf:coverb[ba]:b_dprime:sag[central] | 0.10 | 0.19 | 0.49 |
| scale(Theta):iaf:type[noun]:b_dprime:sag[anterior] | -0.23 | 0.19 | -1.24 |
| scale(Theta):iaf:type[noun]:b_dprime:sag[central] | 0.12 | 0.19 | 0.62 |
| scale(Theta):coverb[ba]:type[noun]:b_dprime:sag[anterior] | -1.96 | 1.81 | -1.08 |
| scale(Theta):coverb[ba]:type[noun]:b_dprime:sag[central] | 1.37 | 1.85 | 0.74 |
| iaf:coverb[ba]:type[noun]:b_dprime:sag[anterior] | 0.08 | 0.14 | 0.59 |
| iaf:coverb[ba]:type[noun]:b_dprime:sag[central] | -0.06 | 0.15 | -0.37 |
| scale(Theta):iaf:coverb[ba]:type[noun]:lat[left] | 0.25 | 0.15 | 1.68 |
| scale(Theta):iaf:coverb[ba]:type[noun]:lat[midline] | -0.34 | 0.12 | -2.76 |
| scale(Theta):iaf:coverb[ba]:b_dprime:lat[left] | 0.15 | 0.22 | 0.68 |
| scale(Theta):iaf:coverb[ba]:b_dprime:lat[midline] | -0.19 | 0.18 | -1.09 |
| scale(Theta):iaf:type[noun]:b_dprime:lat[left] | 0.24 | 0.22 | 1.09 |
| scale(Theta):iaf:type[noun]:b_dprime:lat[midline] | -0.29 | 0.17 | -1.70 |
| scale(Theta):coverb[ba]:type[noun]:b_dprime:lat[left] | 2.55 | 2.09 | 1.22 |
| scale(Theta):coverb[ba]:type[noun]:b_dprime:lat[midline] | -3.35 | 1.66 | -2.02 |
| iaf:coverb[ba]:type[noun]:b_dprime:lat[left] | -0.06 | 0.14 | -0.47 |
| iaf:coverb[ba]:type[noun]:b_dprime:lat[midline] | 0.05 | 0.14 | 0.38 |
| scale(Theta):iaf:coverb[ba]:sag[anterior]:lat[left] | -0.05 | 0.20 | -0.27 |
| scale(Theta):iaf:coverb[ba]:sag[central]:lat[left] | 0.07 | 0.20 | 0.36 |
| scale(Theta):iaf:coverb[ba]:sag[anterior]:lat[midline] | -0.13 | 0.18 | -0.72 |
| scale(Theta):iaf:coverb[ba]:sag[central]:lat[midline] | 0.04 | 0.17 | 0.23 |
| scale(Theta):iaf:type[noun]:sag[anterior]:lat[left] | -0.09 | 0.20 | -0.47 |
| scale(Theta):iaf:type[noun]:sag[central]:lat[left] | 0.14 | 0.20 | 0.70 |
| scale(Theta):iaf:type[noun]:sag[anterior]:lat[midline] | -0.18 | 0.18 | -0.99 |
| scale(Theta):iaf:type[noun]:sag[central]:lat[midline] | 0.05 | 0.17 | 0.30 |

Table 6: Model output for REM theta model *(continued)*

| | Estimate | Std. Error | t value |
|---|---|---|---|
| scale(Theta):coverb[ba]:type[noun]:sag[anterior]:lat[left] | 0.05 | 1.84 | 0.03 |
| scale(Theta):coverb[ba]:type[noun]:sag[central]:lat[left] | 1.56 | 1.89 | 0.83 |
| scale(Theta):coverb[ba]:type[noun]:sag[anterior]:lat[midline] | -0.59 | 1.63 | -0.36 |
| scale(Theta):coverb[ba]:type[noun]:sag[central]:lat[midline] | -0.74 | 1.61 | -0.46 |
| iaf:coverb[ba]:type[noun]:sag[anterior]:lat[left] | 0.09 | 0.14 | 0.65 |
| iaf:coverb[ba]:type[noun]:sag[central]:lat[left] | -0.07 | 0.13 | -0.56 |
| iaf:coverb[ba]:type[noun]:sag[anterior]:lat[midline] | -0.04 | 0.11 | -0.38 |
| iaf:coverb[ba]:type[noun]:sag[central]:lat[midline] | 0.05 | 0.12 | 0.42 |
| scale(Theta):iaf:b_dprime:sag[anterior]:lat[left] | -0.12 | 0.28 | -0.43 |
| scale(Theta):iaf:b_dprime:sag[central]:lat[left] | 0.15 | 0.28 | 0.53 |
| scale(Theta):iaf:b_dprime:sag[anterior]:lat[midline] | 0.07 | 0.24 | 0.30 |
| scale(Theta):iaf:b_dprime:sag[central]:lat[midline] | -0.10 | 0.24 | -0.43 |
| scale(Theta):coverb[ba]:b_dprime:sag[anterior]:lat[left] | 0.20 | 2.68 | 0.07 |
| scale(Theta):coverb[ba]:b_dprime:sag[central]:lat[left] | -0.29 | 2.74 | -0.11 |
| scale(Theta):coverb[ba]:b_dprime:sag[anterior]:lat[midline] | -1.29 | 2.30 | -0.56 |
| scale(Theta):coverb[ba]:b_dprime:sag[central]:lat[midline] | 0.77 | 2.35 | 0.33 |
| iaf:coverb[ba]:b_dprime:sag[anterior]:lat[left] | 0.11 | 0.19 | 0.57 |
| iaf:coverb[ba]:b_dprime:sag[central]:lat[left] | -0.05 | 0.20 | -0.27 |
| iaf:coverb[ba]:b_dprime:sag[anterior]:lat[midline] | 0.01 | 0.20 | 0.04 |
| iaf:coverb[ba]:b_dprime:sag[central]:lat[midline] | 0.08 | 0.24 | 0.33 |
| scale(Theta):type[noun]:b_dprime:sag[anterior]:lat[left] | -1.10 | 2.67 | -0.41 |
| scale(Theta):type[noun]:b_dprime:sag[central]:lat[left] | 2.05 | 2.74 | 0.75 |
| scale(Theta):type[noun]:b_dprime:sag[anterior]:lat[midline] | -0.46 | 2.29 | -0.20 |
| scale(Theta):type[noun]:b_dprime:sag[central]:lat[midline] | -1.31 | 2.33 | -0.56 |
| iaf:type[noun]:b_dprime:sag[anterior]:lat[left] | 0.05 | 0.19 | 0.28 |
| iaf:type[noun]:b_dprime:sag[central]:lat[left] | 0.02 | 0.20 | 0.11 |
| iaf:type[noun]:b_dprime:sag[anterior]:lat[midline] | 0.15 | 0.20 | 0.77 |
| iaf:type[noun]:b_dprime:sag[central]:lat[midline] | -0.11 | 0.23 | -0.47 |
| coverb[ba]:type[noun]:b_dprime:sag[anterior]:lat[left] | 0.42 | 1.82 | 0.23 |
| coverb[ba]:type[noun]:b_dprime:sag[central]:lat[left] | -1.57 | 1.96 | -0.80 |
| coverb[ba]:type[noun]:b_dprime:sag[anterior]:lat[midline] | -0.97 | 1.93 | -0.50 |
| coverb[ba]:type[noun]:b_dprime:sag[central]:lat[midline] | 2.35 | 2.34 | 1.01 |
| scale(Theta):iaf:coverb[ba]:type[noun]:b_dprime:sag[anterior] | 0.20 | 0.19 | 1.08 |
| scale(Theta):iaf:coverb[ba]:type[noun]:b_dprime:sag[central] | -0.15 | 0.19 | -0.79 |
| scale(Theta):iaf:coverb[ba]:type[noun]:b_dprime:lat[left] | -0.28 | 0.22 | -1.28 |
| scale(Theta):iaf:coverb[ba]:type[noun]:b_dprime:lat[midline] | 0.36 | 0.17 | 2.07 |
| scale(Theta):iaf:coverb[ba]:type[noun]:sag[anterior]:lat[left] | 0.02 | 0.20 | 0.10 |
| scale(Theta):iaf:coverb[ba]:type[noun]:sag[central]:lat[left] | -0.18 | 0.20 | -0.87 |
| scale(Theta):iaf:coverb[ba]:type[noun]:sag[anterior]:lat[midline] | 0.05 | 0.18 | 0.27 |
| scale(Theta):iaf:coverb[ba]:type[noun]:sag[central]:lat[midline] | 0.09 | 0.17 | 0.50 |
| scale(Theta):iaf:coverb[ba]:b_dprime:sag[anterior]:lat[left] | -0.01 | 0.28 | -0.05 |
| scale(Theta):iaf:coverb[ba]:b_dprime:sag[central]:lat[left] | 0.03 | 0.28 | 0.09 |
| scale(Theta):iaf:coverb[ba]:b_dprime:sag[anterior]:lat[midline] | 0.13 | 0.24 | 0.55 |
| scale(Theta):iaf:coverb[ba]:b_dprime:sag[central]:lat[midline] | -0.08 | 0.24 | -0.31 |
| scale(Theta):iaf:type[noun]:b_dprime:sag[anterior]:lat[left] | 0.12 | 0.28 | 0.44 |
| scale(Theta):iaf:type[noun]:b_dprime:sag[central]:lat[left] | -0.21 | 0.28 | -0.76 |
| scale(Theta):iaf:type[noun]:b_dprime:sag[anterior]:lat[midline] | 0.05 | 0.24 | 0.22 |
| scale(Theta):iaf:type[noun]:b_dprime:sag[central]:lat[midline] | 0.13 | 0.24 | 0.55 |
| scale(Theta):coverb[ba]:type[noun]:b_dprime:sag[anterior]:lat[left] | -0.68 | 2.68 | -0.26 |
| scale(Theta):coverb[ba]:type[noun]:b_dprime:sag[central]:lat[left] | -1.45 | 2.74 | -0.53 |
| scale(Theta):coverb[ba]:type[noun]:b_dprime:sag[anterior]:lat[midline] | 1.99 | 2.30 | 0.87 |
| scale(Theta):coverb[ba]:type[noun]:b_dprime:sag[central]:lat[midline] | 0.10 | 2.33 | 0.04 |
| iaf:coverb[ba]:type[noun]:b_dprime:sag[anterior]:lat[left] | -0.04 | 0.19 | -0.22 |
| iaf:coverb[ba]:type[noun]:b_dprime:sag[central]:lat[left] | 0.16 | 0.20 | 0.82 |
| iaf:coverb[ba]:type[noun]:b_dprime:sag[anterior]:lat[midline] | 0.10 | 0.20 | 0.52 |
| iaf:coverb[ba]:type[noun]:b_dprime:sag[central]:lat[midline] | -0.24 | 0.23 | -1.04 |
| scale(Theta):iaf:coverb[ba]:type[noun]:b_dprime:sag[anterior]:lat[left] | 0.06 | 0.28 | 0.21 |

Table 6: Model output for REM theta model *(continued)*

| | Estimate | Std. Error | t value |
|---|---|---|---|
| scale(Theta):iaf:coverb[ba]:type[noun]:b_dprime:sag[central]:lat[left] | 0.16 | 0.28 | 0.57 |
| scale(Theta):iaf:coverb[ba]:type[noun]:b_dprime:sag[anterior]:lat[midline] | -0.20 | 0.24 | -0.85 |
| scale(Theta):iaf:coverb[ba]:type[noun]:b_dprime:sag[central]:lat[midline] | -0.02 | 0.24 | -0.08 |

Appendix A.4. Model summary and Wald Test output for time spent in REM model

Table 7: Wald Test output for time spent in REM model

| | Chisq | Df | Pr(>Chisq) |
|---|---|---|---|
| scale(REMperc) | 0.02 | 1 | 0.89 |
| iaf | 1.80 | 1 | 0.18 |
| coverb | 2.85 | 1 | 0.09 |
| type | 4.43 | 1 | 0.04 |
| b_dprime | 38.20 | 1 | 0.00 |
| scale(REMperc):iaf | 0.10 | 1 | 0.75 |
| scale(REMperc):coverb | 3.40 | 1 | 0.07 |
| iaf:coverb | 17.75 | 1 | 0.00 |
| scale(REMperc):type | 0.00 | 1 | 0.95 |
| iaf:type | 2.26 | 1 | 0.13 |
| coverb:type | 8.08 | 1 | 0.00 |
| scale(REMperc):b_dprime | 0.52 | 1 | 0.47 |
| iaf:b_dprime | 5.45 | 1 | 0.02 |
| coverb:b_dprime | 1.14 | 1 | 0.29 |
| type:b_dprime | 0.05 | 1 | 0.83 |
| scale(REMperc):iaf:coverb | 1.71 | 1 | 0.19 |
| scale(REMperc):iaf:type | 0.42 | 1 | 0.52 |
| scale(REMperc):coverb:type | 7.41 | 1 | 0.01 |
| iaf:coverb:type | 12.58 | 1 | 0.00 |
| scale(REMperc):iaf:b_dprime | 1.39 | 1 | 0.24 |
| scale(REMperc):coverb:b_dprime | 4.34 | 1 | 0.04 |
| iaf:coverb:b_dprime | 10.66 | 1 | 0.00 |
| scale(REMperc):type:b_dprime | 0.28 | 1 | 0.60 |
| iaf:type:b_dprime | 1.70 | 1 | 0.19 |
| coverb:type:b_dprime | 0.86 | 1 | 0.35 |
| scale(REMperc):iaf:coverb:type | 0.15 | 1 | 0.70 |
| scale(REMperc):iaf:coverb:b_dprime | 0.69 | 1 | 0.41 |
| scale(REMperc):iaf:type:b_dprime | 0.01 | 1 | 0.94 |
| scale(REMperc):coverb:type:b_dprime | 0.38 | 1 | 0.54 |
| iaf:coverb:type:b_dprime | 0.33 | 1 | 0.56 |
| scale(REMperc):iaf:coverb:type:b_dprime | 0.40 | 1 | 0.53 |

Table 8: Model output for time spent in REM model

| | Estimate | Std. Error | t value |
|---|---|---|---|
| (Intercept) | 2.41 | 3.41 | 0.70 |
| scale(REMperc) | -4.21 | 4.13 | -1.02 |
| iaf | -0.27 | 0.33 | -0.80 |
| coverb[ba] | -7.23 | 1.89 | -3.83 |
| type[noun] | -1.22 | 1.46 | -0.84 |
| b_dprime | 1.98 | 2.67 | 0.74 |
| scale(REMperc):iaf | 0.39 | 0.40 | 0.98 |
| scale(REMperc):coverb[ba] | -3.45 | 1.97 | -1.75 |
| iaf:coverb[ba] | 0.74 | 0.18 | 4.10 |
| scale(REMperc):type[noun] | 0.44 | 1.55 | 0.28 |
| iaf:type[noun] | 0.13 | 0.14 | 0.93 |
| coverb[ba]:type[noun] | 3.01 | 1.27 | 2.37 |
| scale(REMperc):b_dprime | 5.71 | 4.25 | 1.35 |
| iaf:b_dprime | -0.06 | 0.26 | -0.24 |
| coverb[ba]:b_dprime | 8.24 | 2.56 | 3.22 |
| type[noun]:b_dprime | 2.30 | 1.76 | 1.31 |
| scale(REMperc):iaf:coverb[ba] | 0.30 | 0.19 | 1.58 |
| scale(REMperc):iaf:type[noun] | -0.04 | 0.15 | -0.25 |
| scale(REMperc):coverb[ba]:type[noun] | -0.69 | 1.59 | -0.44 |
| iaf:coverb[ba]:type[noun] | -0.32 | 0.12 | -2.59 |
| scale(REMperc):iaf:b_dprime | -0.55 | 0.42 | -1.32 |
| scale(REMperc):coverb[ba]:b_dprime | 3.73 | 3.53 | 1.06 |
| iaf:coverb[ba]:b_dprime | -0.86 | 0.27 | -3.23 |
| scale(REMperc):type[noun]:b_dprime | 1.76 | 3.32 | 0.53 |
| iaf:type[noun]:b_dprime | -0.24 | 0.18 | -1.34 |
| coverb[ba]:type[noun]:b_dprime | 1.16 | 1.92 | 0.61 |
| scale(REMperc):iaf:coverb[ba]:type[noun] | 0.11 | 0.16 | 0.72 |
| scale(REMperc):iaf:coverb[ba]:b_dprime | -0.35 | 0.36 | -0.96 |
| scale(REMperc):iaf:type[noun]:b_dprime | -0.18 | 0.34 | -0.53 |
| scale(REMperc):coverb[ba]:type[noun]:b_dprime | 2.43 | 4.07 | 0.60 |
| iaf:coverb[ba]:type[noun]:b_dprime | -0.13 | 0.20 | -0.68 |
| scale(REMperc):iaf:coverb[ba]:type[noun]:b_dprime | -0.27 | 0.42 | -0.63 |

Appendix B. Visualisations of significant interactions and pairwise comparisons between variables of interest

Figure 1: Significant interactions in the behavioural and PAC models

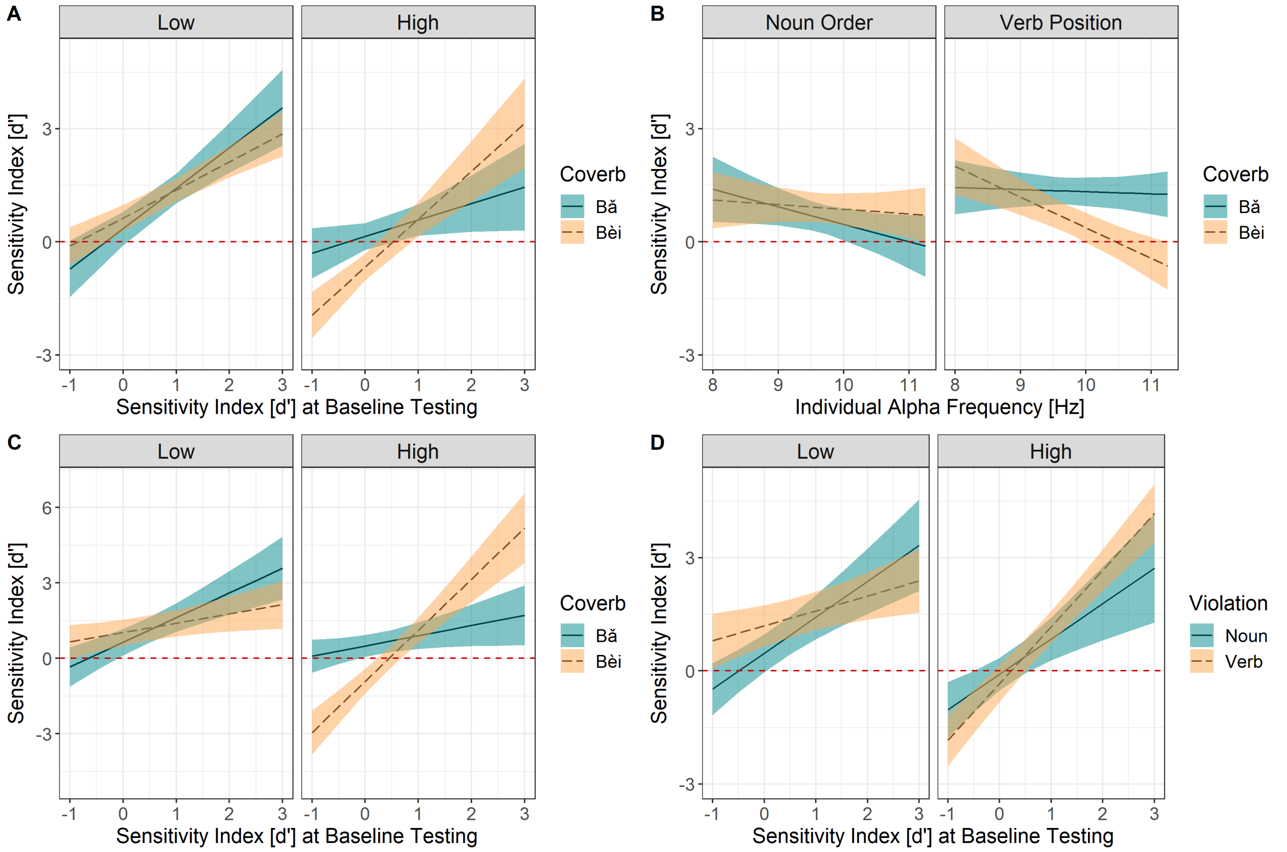


**Note:** All interactions are predicting performance scores at delayed testing (on the y-axis) quantified using d-prime (fitted model values). For all plots, the shaded areas of colour represent 83% confidence intervals and the red dashed line intercepting the y-axis at 0 represents chance performance (d' = 0). **(A)** IAF x Coverb x Baseline d' interaction from the behavioural model. Baseline performance values (d') are depicted on the x-axis, from lower performance values (left) to higher performance values (right), with higher values reflecting higher performance at baseline testing. Additionally, plot is faceted by IAF on the top axis, split into 2 facets, with the left facet showing the relationship between baseline performance and delayed d-prime for lower IAF individuals and the right facet showing this relationship for higher IAF individuals. IAF was dichotomised on this plot for visualisation purposes only. **(B)** IAF x Coverb x Type interaction from the PAC model. IAF values are shown on the x-axis, from lower IAF values (left) to higher IAF values (right). Violation Type is shown on the top axis, split into 2 facets, with noun order violations on the left and verb positioning violations on the right. **(C)** IAF x Coverb x Baseline d' interaction from the PAC model. Baseline performance values (d') are depicted on the x-axis, from lower performance values (left) to higher performance values (right), with higher values reflecting higher performance at baseline testing. IAF is shown on the top axis, split into 2 facets, with the left facet showing the relationship between baseline performance and delayed d-prime for lower IAF individuals and the right facet showing this relationship for higher IAF individuals. IAF was dichotomised on this plot for visualisation purposes only. **(D)** IAF x Violation Type x Baseline d' interaction from the PAC model. Baseline performance values (d') are depicted on the x-axis, from lower performance values (left) to higher performance values (right), with higher values reflecting higher performance at baseline testing. IAF is shown on the top axis, split into 2 facets, with the left facet showing the relationship between baseline performance and delayed d-prime for lower IAF individuals and the right facet showing this relationship for higher IAF individuals. IAF was dichotomised on this plot for visualisation purposes only.

Figure 2: Significant interactions in the REM theta and proportion models

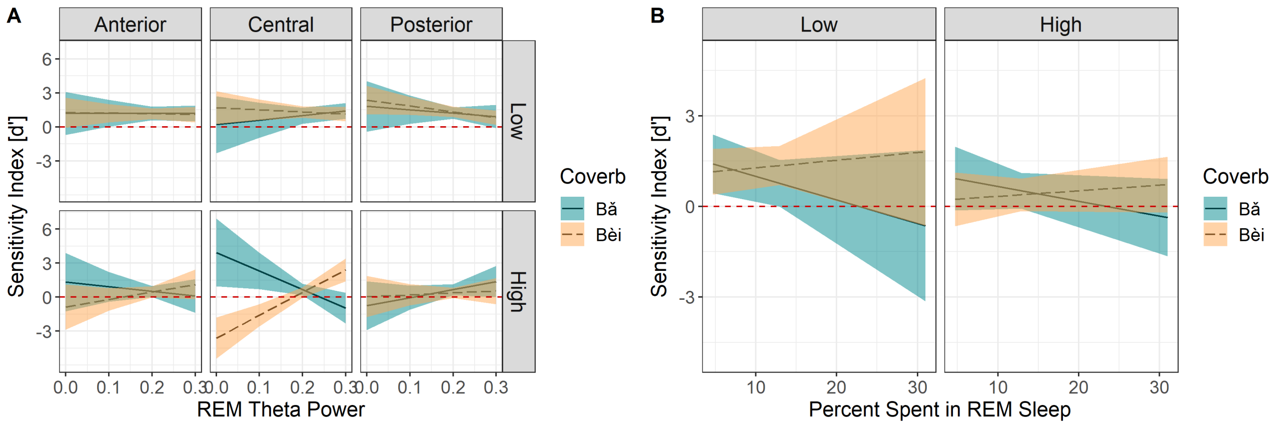


**Note:** All interactions are predicting performance scores at delayed testing (on the y-axis) quantified using d-prime (fitted model values). For all plots, the shaded areas of colour represent 83% confidence intervals and the red dashed line intercepting the y-axis at 0 represents chance performance (d' = 0). **(A)** REM Theta x IAF x Coverb x Saggitality interaction from the REM theta power model. REM Theta Power is shown on the x-axis, from lower power values (left) to higher power values (right), with higher power values reflecting higher power spectral density values. Saggitality is shown on the top axis, split into 3 facets, moving from anterior (left) to central (middle) and posterior (right), depicting topographical positioning. Additionally, this plot is faceted by IAF on the right axis, split into 2 facets, with the left facet showing the relationship between baseline performance and delayed d-prime for lower IAF individuals and the right facet showing this relationship for higher IAF individuals. IAF was dichotomised on this plot for visualisation purposes only. **(B)** Percentage of REM x Coverb x Violation Type interaction from the percentage of REM model. Percentage of time spent in REM sleep is shown along the x-axis, from lower values (left) to higher values (right), with higher values depicting higher percentage of time spent in REM. IAF is shown on the top axis, split into 2 facets, with the left facet showing the relationship between baseline performance and delayed d-prime for lower IAF individuals and the right facet showing this relationship for higher IAF individuals. IAF was dichotomised on this plot for visualisation purposes only.

Figure 3: Correlogram of pairwise comparisons between variables of interest

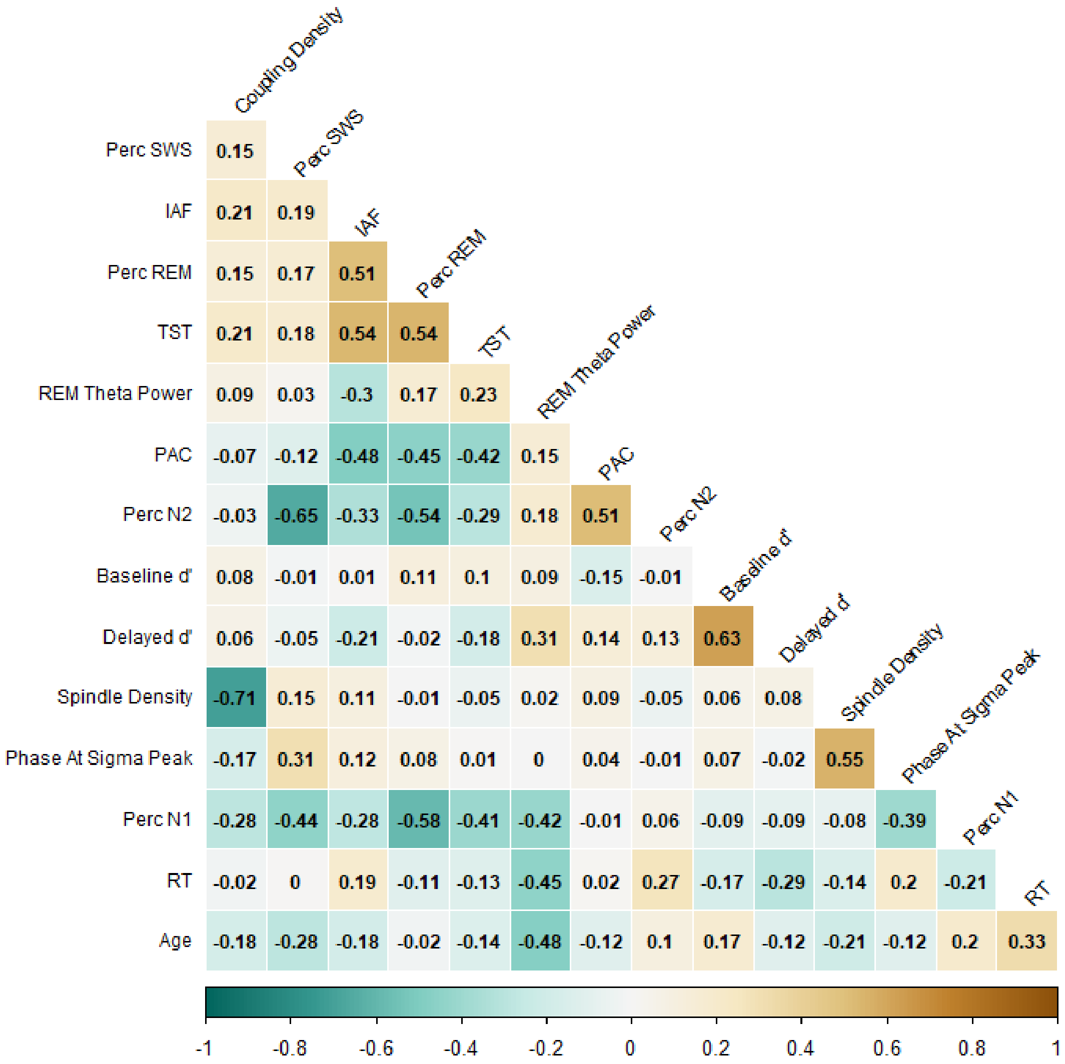


**Note:** Strength and direction of the relationship between variables is shown on along the bottom, with negative relationships being shown in blue, and positive relationships being shown in tan. Additionally, the strength of the relationship is shown via colour opacity, with darker colours indicating a stronger relationship. Significance values (p) are shown in the corresponding box of each relationship.

Figure 4: Correlation between SO-spindle coupling strength and Individual Alpha Frequency during stage 2 and stage 3 sleep

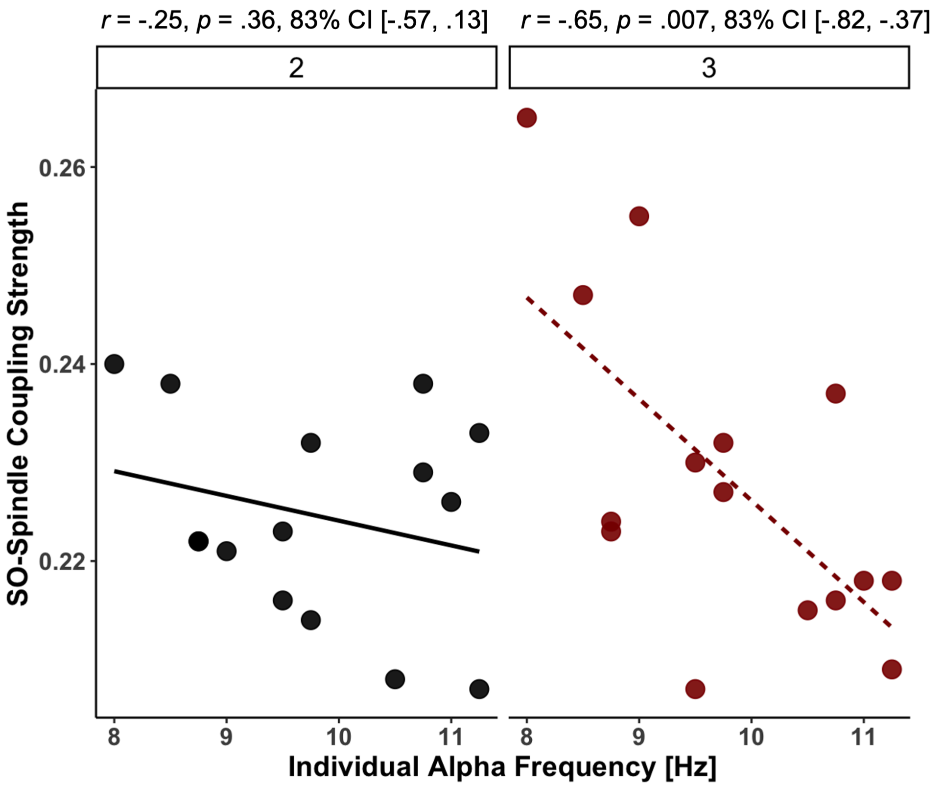


**Note:** PAC values are shown on the y-axis, from lower coupling values (bottom) to higher coupling values (top), with higher coupling values indicating better coupling strength between slow-oscillations (SOs) and spindles. IAF values are shown on the x-axis, from lower IAF values (left) to higher IAF values (right). Sleep stage is shown along the the top axis, split into 2 facets, with the left showing the relationship between PAC and IAF during stage 2 sleep and the right showing this relationship during stage 3 sleep. Statistical information for each relationship are shown above each facet.

Appendix C. Raw scores for hits and false alarms per subject per condition

Table 9: Raw hit and false alarms per subject and sentence type

| Subject | Session | Sentence Type | Coverb | Hits | False Alarms |
|---|---|---|---|---|---|
| CP01 | Baseline | noun | ba | 22 | 11 |
| CP01 | Delayed | noun | ba | 18 | 8 |
| CP01 | Baseline | noun | bei | 18 | 17 |
| CP01 | Delayed | noun | bei | 18 | 12 |
| CP01 | Baseline | verb | ba | 22 | 8 |
| CP01 | Delayed | verb | ba | 18 | 12 |
| CP01 | Baseline | verb | bei | 18 | 16 |
| CP01 | Delayed | verb | bei | 18 | 14 |
| CP02 | Baseline | noun | ba | 25 | 14 |
| CP02 | Delayed | noun | ba | 32 | 20 |
| CP02 | Baseline | noun | bei | 27 | 17 |
| CP02 | Delayed | noun | bei | 30 | 18 |
| CP02 | Baseline | verb | ba | 25 | 12 |
| CP02 | Delayed | verb | ba | 32 | 18 |
| CP02 | Baseline | verb | bei | 27 | 12 |
| CP02 | Delayed | verb | bei | 30 | 18 |
| CP03 | Baseline | noun | ba | 25 | 22 |
| CP03 | Delayed | noun | ba | 22 | 13 |
| CP03 | Baseline | noun | bei | 28 | 20 |
| CP03 | Delayed | noun | bei | 26 | 22 |
| CP03 | Baseline | verb | ba | 25 | 8 |
| CP03 | Delayed | verb | ba | 22 | 4 |
| CP03 | Baseline | verb | bei | 28 | 10 |
| CP03 | Delayed | verb | bei | 26 | 6 |
| CP04 | Baseline | noun | ba | 16 | 11 |
| CP04 | Delayed | noun | ba | 16 | 11 |
| CP04 | Baseline | noun | bei | 16 | 9 |
| CP04 | Delayed | noun | bei | 13 | 9 |
| CP04 | Baseline | verb | ba | 16 | 2 |
| CP04 | Delayed | verb | ba | 16 | 14 |
| CP04 | Baseline | verb | bei | 16 | 8 |
| CP04 | Delayed | verb | bei | 13 | 14 |
| CP05 | Baseline | noun | ba | 14 | 14 |
| CP05 | Delayed | noun | ba | 20 | 15 |
| CP05 | Baseline | noun | bei | 23 | 11 |
| CP05 | Delayed | noun | bei | 21 | 16 |
| CP05 | Baseline | verb | ba | 14 | 12 |
| CP05 | Delayed | verb | ba | 20 | 14 |
| CP05 | Baseline | verb | bei | 23 | 4 |
| CP05 | Delayed | verb | bei | 21 | 14 |
| CP06 | Baseline | noun | ba | 21 | 14 |
| CP06 | Delayed | noun | ba | 22 | 12 |
| CP06 | Baseline | noun | bei | 24 | 14 |
| CP06 | Delayed | noun | bei | 16 | 12 |
| CP06 | Baseline | verb | ba | 21 | 12 |
| CP06 | Delayed | verb | ba | 22 | 10 |
| CP06 | Baseline | verb | bei | 24 | 6 |
| CP06 | Delayed | verb | bei | 16 | 8 |
| CP07 | Baseline | noun | ba | 15 | 12 |
| CP07 | Delayed | noun | ba | 3 | 8 |
| CP07 | Baseline | noun | bei | 17 | 8 |
| CP07 | Delayed | noun | bei | 21 | 10 |
| CP07 | Baseline | verb | ba | 15 | 10 |
| CP07 | Delayed | verb | ba | 3 | 12 |
| CP07 | Baseline | verb | bei | 17 | 6 |

Table 9: Raw hit and false alarms per subject and sentence type *(continued)*

| Subject | Session | Sentence Type | Coverb | Hits | False Alarms |
|---|---|---|---|---|---|
| CP07 | Delayed | verb | bei | 21 | 18 |
| CP09 | Baseline | noun | ba | 27 | 20 |
| CP09 | Delayed | noun | ba | 21 | 19 |
| CP09 | Baseline | noun | bei | 27 | 13 |
| CP09 | Delayed | noun | bei | 26 | 17 |
| CP09 | Baseline | verb | ba | 27 | 10 |
| CP09 | Delayed | verb | ba | 21 | 10 |
| CP09 | Baseline | verb | bei | 27 | 12 |
| CP09 | Delayed | verb | bei | 26 | 18 |
| CP10 | Baseline | noun | ba | 32 | 12 |
| CP10 | Delayed | noun | ba | 35 | 6 |
| CP10 | Baseline | noun | bei | 32 | 6 |
| CP10 | Delayed | noun | bei | 35 | 4 |
| CP10 | Baseline | verb | ba | 32 | 2 |
| CP10 | Delayed | verb | ba | 35 | 1 |
| CP10 | Baseline | verb | bei | 32 | 2 |
| CP10 | Delayed | verb | bei | 35 | 1 |
| CP11 | Baseline | noun | ba | 23 | 15 |
| CP11 | Delayed | noun | ba | 12 | 9 |
| CP11 | Baseline | noun | bei | 33 | 20 |
| CP11 | Delayed | noun | bei | 22 | 9 |
| CP11 | Baseline | verb | ba | 23 | 6 |
| CP11 | Delayed | verb | ba | 12 | 4 |
| CP11 | Baseline | verb | bei | 33 | 1 |
| CP11 | Delayed | verb | bei | 22 | 1 |
| CP12 | Baseline | noun | ba | 17 | 12 |
| CP12 | Delayed | noun | ba | 23 | 15 |
| CP12 | Baseline | noun | bei | 30 | 13 |
| CP12 | Delayed | noun | bei | 35 | 20 |
| CP12 | Baseline | verb | ba | 17 | 2 |
| CP12 | Delayed | verb | ba | 23 | 4 |
| CP12 | Baseline | verb | bei | 30 | 1 |
| CP12 | Delayed | verb | bei | 35 | 1 |
| CP13 | Baseline | noun | ba | 27 | 13 |
| CP13 | Delayed | noun | ba | 28 | 12 |
| CP13 | Baseline | noun | bei | 26 | 15 |
| CP13 | Delayed | noun | bei | 35 | 23 |
| CP13 | Baseline | verb | ba | 27 | 4 |
| CP13 | Delayed | verb | ba | 28 | 1 |
| CP13 | Baseline | verb | bei | 26 | 8 |
| CP13 | Delayed | verb | bei | 35 | 1 |
| CP14 | Baseline | noun | ba | 27 | 18 |
| CP14 | Delayed | noun | ba | 35 | 23 |
| CP14 | Baseline | noun | bei | 34 | 18 |
| CP14 | Delayed | noun | bei | 35 | 14 |
| CP14 | Baseline | verb | ba | 27 | 4 |
| CP14 | Delayed | verb | ba | 35 | 2 |
| CP14 | Baseline | verb | bei | 34 | 6 |
| CP14 | Delayed | verb | bei | 35 | 4 |
| CP15 | Baseline | noun | ba | 32 | 12 |
| CP15 | Delayed | noun | ba | 26 | 6 |
| CP15 | Baseline | noun | bei | 18 | 12 |
| CP15 | Delayed | noun | bei | 18 | 8 |
| CP15 | Baseline | verb | ba | 32 | 2 |
| CP15 | Delayed | verb | ba | 26 | 4 |
| CP15 | Baseline | verb | bei | 18 | 4 |
| CP15 | Delayed | verb | bei | 18 | 1 |

Table 9: Raw hit and false alarms per subject and sentence type *(continued)*

| Subject | Session | Sentence Type | Coverb | Hits | False Alarms |
|---|---|---|---|---|---|
| CP16 | Baseline | noun | ba | 24 | 15 |
| CP16 | Delayed | noun | ba | 16 | 23 |
| CP16 | Baseline | noun | bei | 23 | 13 |
| CP16 | Delayed | noun | bei | 14 | 21 |
| CP16 | Baseline | verb | ba | 24 | 2 |
| CP16 | Delayed | verb | ba | 16 | 2 |
| CP16 | Baseline | verb | bei | 23 | 4 |
| CP16 | Delayed | verb | bei | 14 | 2 |
| CP17 | Baseline | noun | ba | 3 | 3 |
| CP17 | Delayed | noun | ba | 6 | 4 |
| CP17 | Baseline | noun | bei | 4 | 4 |
| CP17 | Delayed | noun | bei | 7 | 7 |
| CP17 | Baseline | verb | ba | 3 | 2 |
| CP17 | Delayed | verb | ba | 6 | 4 |
| CP17 | Baseline | verb | bei | 4 | 1 |
| CP17 | Delayed | verb | bei | 7 | 6 |
| CP18 | Baseline | noun | ba | 27 | 15 |
| CP18 | Delayed | noun | ba | 26 | 15 |
| CP18 | Baseline | noun | bei | 26 | 22 |
| CP18 | Delayed | noun | bei | 27 | 21 |
| CP18 | Baseline | verb | ba | 27 | 22 |
| CP18 | Delayed | verb | ba | 26 | 14 |
| CP18 | Baseline | verb | bei | 26 | 18 |
| CP18 | Delayed | verb | bei | 27 | 18 |
| CP19 | Baseline | noun | ba | 30 | 19 |
| CP19 | Delayed | noun | ba | 13 | 16 |
| CP19 | Baseline | noun | bei | 25 | 15 |
| CP19 | Delayed | noun | bei | 23 | 22 |
| CP19 | Baseline | verb | ba | 30 | 1 |
| CP19 | Delayed | verb | ba | 13 | 1 |
| CP19 | Baseline | verb | bei | 25 | 2 |
| CP19 | Delayed | verb | bei | 23 | 2 |
| CP20 | Baseline | noun | ba | 23 | 11 |
| CP20 | Delayed | noun | ba | 12 | 5 |
| CP20 | Baseline | noun | bei | 17 | 13 |
| CP20 | Delayed | noun | bei | 9 | 1 |
| CP20 | Baseline | verb | ba | 23 | 6 |
| CP20 | Delayed | verb | ba | 12 | 6 |
| CP20 | Baseline | verb | bei | 17 | 14 |
| CP20 | Delayed | verb | bei | 9 | 6 |
| CP21 | Baseline | noun | ba | 27 | 14 |
| CP21 | Delayed | noun | ba | 21 | 12 |
| CP21 | Baseline | noun | bei | 20 | 18 |
| CP21 | Delayed | noun | bei | 14 | 16 |
| CP21 | Baseline | verb | ba | 27 | 10 |
| CP21 | Delayed | verb | ba | 21 | 18 |
| CP21 | Baseline | verb | bei | 20 | 18 |
| CP21 | Delayed | verb | bei | 14 | 22 |
| CP22 | Baseline | noun | ba | 24 | 17 |
| CP22 | Delayed | noun | ba | 30 | 5 |
| CP22 | Baseline | noun | bei | 29 | 7 |
| CP22 | Delayed | noun | bei | 32 | 6 |
| CP22 | Baseline | verb | ba | 24 | 12 |
| CP22 | Delayed | verb | ba | 30 | 1 |
| CP22 | Baseline | verb | bei | 29 | 20 |
| CP22 | Delayed | verb | bei | 32 | 2 |
| CP23 | Baseline | noun | ba | 28 | 20 |

Table 9: Raw hit and false alarms per subject and sentence type *(continued)*

| Subject | Session | Sentence Type | Coverb | Hits | False Alarms |
|---|---|---|---|---|---|
| CP23 | Delayed | noun | ba | 30 | 23 |
| CP23 | Baseline | noun | bei | 31 | 21 |
| CP23 | Delayed | noun | bei | 31 | 22 |
| CP23 | Baseline | verb | ba | 28 | 8 |
| CP23 | Delayed | verb | ba | 30 | 8 |
| CP23 | Baseline | verb | bei | 31 | 14 |
| CP23 | Delayed | verb | bei | 31 | 2 |
| CP24 | Baseline | noun | ba | 21 | 13 |
| CP24 | Delayed | noun | ba | 27 | 15 |
| CP24 | Baseline | noun | bei | 19 | 14 |
| CP24 | Delayed | noun | bei | 21 | 19 |
| CP24 | Baseline | verb | ba | 21 | 8 |
| CP24 | Delayed | verb | ba | 27 | 4 |
| CP24 | Baseline | verb | bei | 19 | 6 |
| CP24 | Delayed | verb | bei | 21 | 6 |
| CP25 | Baseline | noun | ba | 30 | 17 |
| CP25 | Delayed | noun | ba | 35 | 11 |
| CP25 | Baseline | noun | bei | 31 | 17 |
| CP25 | Delayed | noun | bei | 35 | 7 |
| CP25 | Baseline | verb | ba | 30 | 6 |
| CP25 | Delayed | verb | ba | 35 | 1 |
| CP25 | Baseline | verb | bei | 31 | 6 |
| CP25 | Delayed | verb | bei | 35 | 4 |
| CP26 | Baseline | noun | ba | 25 | 16 |
| CP26 | Delayed | noun | ba | 34 | 21 |
| CP26 | Baseline | noun | bei | 22 | 16 |
| CP26 | Delayed | noun | bei | 6 | 7 |
| CP26 | Baseline | verb | ba | 25 | 12 |
| CP26 | Delayed | verb | ba | 34 | 6 |
| CP26 | Baseline | verb | bei | 22 | 12 |
| CP26 | Delayed | verb | bei | 6 | 23 |
| CP27 | Baseline | noun | ba | 28 | 18 |
| CP27 | Delayed | noun | ba | 23 | 20 |
| CP27 | Baseline | noun | bei | 22 | 21 |
| CP27 | Delayed | noun | bei | 29 | 16 |
| CP27 | Baseline | verb | ba | 28 | 1 |
| CP27 | Delayed | verb | ba | 23 | 1 |
| CP27 | Baseline | verb | bei | 22 | 1 |
| CP27 | Delayed | verb | bei | 29 | 1 |
| CP29 | Baseline | noun | ba | 18 | 9 |
| CP29 | Delayed | noun | ba | 19 | 14 |
| CP29 | Baseline | noun | bei | 19 | 9 |
| CP29 | Delayed | noun | bei | 14 | 14 |
| CP29 | Baseline | verb | ba | 18 | 10 |
| CP29 | Delayed | verb | ba | 19 | 14 |
| CP29 | Baseline | verb | bei | 19 | 6 |
| CP29 | Delayed | verb | bei | 14 | 14 |
| CP30 | Baseline | noun | ba | 28 | 21 |
| CP30 | Delayed | noun | ba | 32 | 22 |
| CP30 | Baseline | noun | bei | 25 | 19 |
| CP30 | Delayed | noun | bei | 29 | 23 |
| CP30 | Baseline | verb | ba | 28 | 4 |
| CP30 | Delayed | verb | ba | 32 | 12 |
| CP30 | Baseline | verb | bei | 25 | 6 |
| CP30 | Delayed | verb | bei | 29 | 12 |
| CP31 | Baseline | noun | ba | 25 | 10 |
| CP31 | Delayed | noun | ba | 26 | 16 |

Table 9: Raw hit and false alarms per subject and sentence type *(continued)*

| Subject | Session | Sentence Type | Coverb | Hits | False Alarms |
|---|---|---|---|---|---|
| CP31 | Baseline | noun | bei | 17 | 15 |
| CP31 | Delayed | noun | bei | 25 | 20 |
| CP31 | Baseline | verb | ba | 25 | 6 |
| CP31 | Delayed | verb | ba | 26 | 12 |
| CP31 | Baseline | verb | bei | 17 | 8 |
| CP31 | Delayed | verb | bei | 25 | 10 |
| CP33 | Baseline | noun | ba | 24 | 13 |
| CP33 | Delayed | noun | ba | 13 | 10 |
| CP33 | Baseline | noun | bei | 31 | 17 |
| CP33 | Delayed | noun | bei | 18 | 12 |
| CP33 | Baseline | verb | ba | 24 | 1 |
| CP33 | Delayed | verb | ba | 13 | 1 |
| CP33 | Baseline | verb | bei | 31 | 1 |
| CP33 | Delayed | verb | bei | 18 | 1 |
| CP35 | Baseline | noun | ba | 25 | 18 |
| CP35 | Delayed | noun | ba | 31 | 22 |
| CP35 | Baseline | noun | bei | 27 | 21 |
| CP35 | Delayed | noun | bei | 32 | 21 |
| CP35 | Baseline | verb | ba | 25 | 14 |
| CP35 | Delayed | verb | ba | 31 | 8 |
| CP35 | Baseline | verb | bei | 27 | 18 |
| CP35 | Delayed | verb | bei | 32 | 8 |
| CP36 | Baseline | noun | ba | 34 | 22 |
| CP36 | Delayed | noun | ba | 35 | 23 |
| CP36 | Baseline | noun | bei | 30 | 24 |
| CP36 | Delayed | noun | bei | 35 | 23 |
| CP36 | Baseline | verb | ba | 34 | 6 |
| CP36 | Delayed | verb | ba | 35 | 1 |
| CP36 | Baseline | verb | bei | 30 | 8 |
| CP36 | Delayed | verb | bei | 35 | 1 |
| CP37 | Baseline | noun | ba | 33 | 6 |
| CP37 | Delayed | noun | ba | 35 | 5 |
| CP37 | Baseline | noun | bei | 34 | 11 |
| CP37 | Delayed | noun | bei | 35 | 3 |
| CP37 | Baseline | verb | ba | 33 | 4 |
| CP37 | Delayed | verb | ba | 35 | 1 |
| CP37 | Baseline | verb | bei | 34 | 4 |
| CP37 | Delayed | verb | bei | 35 | 1 |
| CP38 | Baseline | noun | ba | 27 | 13 |
| CP38 | Delayed | noun | ba | 28 | 12 |
| CP38 | Baseline | noun | bei | 18 | 18 |
| CP38 | Delayed | noun | bei | 29 | 19 |
| CP38 | Baseline | verb | ba | 27 | 8 |
| CP38 | Delayed | verb | ba | 28 | 10 |
| CP38 | Baseline | verb | bei | 18 | 12 |
| CP38 | Delayed | verb | bei | 29 | 6 |

Appendix D. Additional analysis outputs

Table 10: n responses missed per condition (Sleep, Wake) and coverb (Bǎ, Bèi)

| Condition | Coverb | n |
|---|---|---|
| Wake | ba | 36 |
| Wake | bei | 33 |
| Sleep | ba | 22 |
| Sleep | bei | 11 |

Note: Table contains the number of responses that were a missed response, which refers to trials in which the maximum response time of 4000ms was exceeded and an incorrect response was recorded.

Table 11: F test to compare two variances

| Estimate | Num df | Denom df | F | p | CI_lower | CI_upper |
|---|---|---|---|---|---|---|
| 1.31 | 71 | 63 | 1.31 | 0.27 | 0.81 | 2.12 |

Table 12: Independant t-test comparing baseline performance between sleep and wake

| Dependent Variable | t | df | p | d | CI_lower | CI_upper |
|---|---|---|---|---|---|---|
| b_dprime | 0.4 | 134 | 0.69 | 0.07 | -0.27 | 0.41 |

Appendix E.1. Link to Study Materials from Cross et al., 2021

The current study is a reanalysis of Cross et al., 202. The stimuli, experimental paradigm and other materials for the original experiment can be found in the following repository:

https://osf.io/f5ksg/?view$_{o}nly = 738200b63e78433eb50e667ea772c08f$

Appendix E.2. Link to Raw Data and Analysis Pipeline Used in the Current Project

All raw data and R scripts created to conduct the analysis for this project, including scripts to calculate d-prime and mixed model analysis, can be found within the open repository linked below:

https://tinyurl.com/turbidus
